# A FUL-SHP-AP2 module regulates fruit development in petunia

**DOI:** 10.64898/2026.03.04.709543

**Authors:** Marian Bemer, Pierre Chambrier, Natali Hernandez Ciro, Patrice Morel, Edelin Roque, Kai Thoris, Eunbae An, Concepción Gómez-Mena, Suzanne Rodrigues Bento, Michiel Vandenbussche

**Affiliations:** Biosystematics group, Wageningen University & Research, Droevendaalsesteeg 1, 6708 PB Wageningen, the Netherlands; Laboratoire de Reproduction et Développement des Plantes, Université de Lyon, ENS de Lyon, UCB Lyon 1, CNRS, INRAE, Lyon 69007, France; Instituto de Biología Molecular y Celular de Plantas (Consejo Superior de Investigaciones Científicas–Universitat Politècnica de València), C/ Ingeniero Fausto Elio s/n, Edif. 8E, 46022 Valencia, Spain

**Keywords:** Fruit development, petunia, pericarp, *FRUITFULL*, *SHATTERPROOF*, *APETALA 2*

## Abstract

Fruit development is a key angiosperm innovation for seed dispersal, yet it remains unclear whether a conserved gene regulatory network operates during early fruit development. We investigated *Petunia × hybrida* homologs of *FRUITFULL* (*FUL*), *SHATTERPROOF* (*SHP*), and *APETALA 2* (*AP2*), transcription factors repeatedly linked to fruit development/ripening. As a model species close to tomato, but producing dry dehiscent fruits like Arabidopsis, petunia is ideal for investigating conserved regulatory mechanisms in early fruit patterning. We used reverse genetics to dissect the roles of the petunia *FUL*, *SHP* and *AP2* homologs in pericarp development. We found that the three petunia *FUL*-like genes redundantly regulate endocarp formation, likely via auxin/brassinosteroid signalling and cell-wall modification. Interestingly, *FBP6* (*SHP-like*) has an opposite effect on endocarp cell morphology, suggesting antagonistic interactions. Mutants in the *AP2-*like *ROB* genes show the most severe pericarp patterning defects, with failure of dehiscence in *rob1/2/3 mutants*. ROB factors repress *FUL*-like activity and, along with FBP6, oppositely regulate auxin/brassinosteroid signalling genes. Our findings uncover a conserved module of antagonistically acting transcription factors (AP2, FUL, SHP co-orthologs) that regulates early pericarp patterning via hormonal signalling, providing a framework for comparative fruit development studies across angiosperms.

**Highlight:** Three key transcription factors in plant reproductive development, FRUITFULL, SHATTERPROOF and APETALA 2, regulate petunia fruit development and together control pericarp patterning in the young fruit.

## Introduction

The evolution of the flower and the fruit enabled the rise of the angiosperms by effectuating efficient pollination and seed dispersal. Decades of comparative research in flower development has revealed that parts of the regulatory network underlying flower development, described by the (A)BC(D)E model (reviewed in, e.g. (Heijmans *et al*., 2012b; Smaczniak, Cezary *et al*., 2012; Bowman & Moyroud, 2024)), are largely conserved throughout the angiosperms, especially the B and C floral organ identity functions. The high diversity of floral forms is partially achieved by shifts in gene expression of these key transcription factors and their targets (for reviews, see e.g. (Moyroud & Glover, 2017; Shan *et al*., 2019). While the conservation of the flower regulatory network has been widely tested, it is still unclear to what extent the fruit regulatory network can be described by a model common to all angiosperms. Interestingly, several flowering genes play an important role in the fruit as well, including members of the MADS-box *APETALA1/FRUITFULL (AP1/FUL)* subfamily and C-class subfamily (including *AGAMOUS* (*AG*) and *SHATTERPROOF 1,2* (*SHP1,2*)), and the *AP2*-like gene *APETALA 2* (*AP2*). This suggests that existing modules have been deployed to facilitate fruit evolution.

Fruit types can be roughly divided into dry fruits, which often disperse their seeds via pod shattering, and fleshy fruits, where the seeds are dispersed by animals. The regulatory network underlying fruit development has been well-studied in the model species *Arabidopsis thaliana* (hereafter: Arabidopsis), which develops dry siliques that form dehiscence zones along which the fruit opens to shatter the seeds. The establishment of cell identities in the silique is regulated by a network of transcription factors, and includes the valve cell-controlling gene *FRUITFULL* (*FUL*) (Gu *et al*., 1998a), the replum identity gene *REPLUMLESS* (*RPL*) (Roeder *et al*., 2003), and the valve margin/dehiscence zone identity genes *SHATTERPROOF 1* (*SHP1*), *SHP2*, *INDEHISCENT* (*IND*) and *ALCATRAZ* (*ALC*) (Liljegren *et al*., 2000; Rajani & Sundaresan, 2001; Liljegren *et al*., 2004) (for review, see (Roeder & Yanofsky, 2006)). Upstream of the replum-and valve margin identity genes, but downstream of FUL, *APETALA 2* (*AP2*) plays an important role in restricting the expression domains of *SHP*, *IND* and *ALC* (Ripoll *et al*., 2011). Within the Brassicaceae family, the gene functions in this regulatory module appear largely conserved (Østergaard *et al*., 2006; Braatz *et al*., 2017; Zhai *et al*., 2019). Outside the Brassicaceae, this appears to be only partially the case, as *IND* is for example a Brassicales-specific bHLH transcription factor (Zumajo-Cardona *et al*., 2025). However, different studies have revealed that orthologs of *FUL*, *SHP* and *AP2* do play a role in tomato fleshy fruit ripening. *Solanum lycopersicum* FUL1 (SlFUL1) and SlFUL2 physically interact with the master ripening regulator RIPENING INHIBITOR (RIN) to initiate aspects of fruit ripening (Bemer *et al*., 2012; Wang *et al*., 2019), while the SHP ortholog TOMATO AGAMOUS-LIKE 1 (TAGL1) also interacts with RIN to induce different ripening aspects (Vrebalov *et al*., 2009). Interestingly, similar to Arabidopsis, the tomato *AP2* co-ortholog *AP2a* plays an antagonistic role and delays fruit ripening (Chung *et al*., 2010; Karlova *et al*., 2011; Wang *et al*., 2019).

While the *FUL*-, *SHP*-and *AP2*-orthologs thus appear important in fruit development/ripening in different species, there is no clear evidence that they act in a conserved network, as studies have focused on fruit dehiscence (Arabidopsis) and fleshy fruit ripening (tomato), two physiologically unrelated processes in distantly related species. To shed more light on a possible conservation of the FUL-SHP-AP2 module, we chose *Petunia x hybrida* (hereafter: petunia) as a model system. Petunia is phylogenetically close to tomato (both Solanaceae family), but produces capsular dry fruits that dehisce to shatter their seeds, similar to Arabidopsis (Pabón-Mora & Litt, 2011). The main difference between the Arabidopsis silique and the petunia capsule, is that in Arabidopsis, the ovules are attached to the ovary wall and a septum divides the silique in two chambers, while the petunia capsule has axile placentation, with the ovules connected to a central placenta (Pabón-Mora & Litt, 2011). In Arabidopsis, the septum connects to the outer carpel walls (the valves) at the replum, and the replum thus separates the two valves, with a few files of valve margin cells developing between valve and replum, forming the dehiscence zone as described above (Roeder, 2006). While a valve margin is not specified in petunia capsular fruits, a few rows of smaller cells do develop at the boundary of the two carpels, which mark the future dehiscence zone, but remain unlignified (Pabón-Mora & Litt, 2011). Because Arabidopsis and petunia show similar fruit developmental trajectories despite their substantial evolutionary distance, petunia provides an ideal model to assess how well the genetic programs underlying dry fruit development are conserved, while its genetic proximity to tomato will facilitate gene regulatory network comparisons underlying dry and fleshy fruit development.

Petunia contains three *AP2* co-orthologs, named *REPRESSOR OF B-FUNCTION 1/2/3 (ROB1/2/3)* and two related *AP2* homologs that belong to the neighbouring *TOE1/2* clade: *BLIND ENHANCER* (*BEN*) and *BROTHER OF BEN* (*BOB*) (Morel *et al*., 2017). It has been shown that during flower development, *ROB1/2/3* and *BEN* redundantly repress the B-function genes in the first whorl, while *BEN* in combination with the *miRNA169 BLIND* (Cartolano *et al*., 2007) restricts the expression of the C-function genes to the inner floral whorls, together illustrating the divergence and molecular diversity of the mechanisms that correspond to the floral A-function. Finally, it was shown that *ROB1/2/3* and *BEN* redundantly, but in an additive manner, negatively regulate the size of the nectaries by preventing their development in the apical region of the ovary (Morel *et al*., 2018). The two petunia, C-class genes are named *PETUNIA MADS BOX GENE 3* (*pMADS3*) and *FLORAL-BINDING PROTEIN 6* (*FBP6*), which are orthologs of Arabidopsis *AGAMOUS* (*AG*) and *SHP1,2* respectively. They fulfil largely overlapping functions in the determination of stamen and carpel identity as well as in floral determinacy (Heijmans *et al*., 2012a), and act as master regulators of nectary development (Morel *et al*., 2018). *FBP6* however, does play a unique role in stigma and style development (Heijmans *et al*., 2012a). The petunia *AP1/FUL* subfamily contains four genes divided over three conserved subclades: *Petunia hybrida* eu*AP1* (*PheuAP1, AP1-clade*), *FLORAL BINDING PROTEIN 26* (*FBP26,* euFULI-clade), *PETUNIA FLOWERING GENE* (*PFG,* euFULI-clade), and *FBP29* (euFULII-clade) (Maheepala *et al*., 2019; Morel *et al*., 2019). These genes are redundantly required for cymose inflorescence development (Morel *et al*., 2019). Quadruple mutants show the most severe phenotype, producing a solitary flower after the production of numerous leaf-like organs on the inflorescence. In addition, it was shown that they repress the B-function genes in the first whorl during flower development (Morel *et al*., 2019). In conclusion, the petunia *AP1/FUL*, AG/*SHP*-and *AP2*-like genes have shown to be important for the regulation of flowering, but their role in fruit development remained to be studied.

Here, we investigated the roles of the petunia *AP1/FUL*, *AP2* and *AG/SHP* co-orthologs during petunia capsular fruit development. We show that the FUL-like transcription factors PFG, FBP26 and FBP29 have redundant roles in early pericarp patterning, the differentiation of the endocarp layers, and endocarp functioning. They influence auxin and brassinosteroid signalling by activating INDOLEACETIC ACID-INDUCED PROTEIN encoding genes (*IAAs*) and transcription factor encoding genes closely homologous to *BRASSINAZOLE-RESISTANT 1* (*BZR1*-like). Additionally, they regulate cell wall modifying genes, including close homologs of *SHOU4*, *TOUCH4* (*TCH4)*, and *IRREGULAR XYLEM 15-LIKE* (*IRX15L).* Interestingly, *PhFUL* gene expression levels are repressed by the AP2-like transcription factors ROB1/2/3 and the MADS-box AG/SHP-clade factor FBP6, which are also required for proper pericarp differentiation. Our mutant analysis suggests that the functions of *ROB1/2/3* and *FBP6* are opposite to those of the *PhFULs*, as *rob1/2/3* and *fbp6* mutants have smaller, regularly-shaped endocarp cells, whereas *phful* mutants have larger and rounder cells compared to WT. While the PhFULs activate *IAA-*and *BZR*-like genes, ROB1/2/3 and FBP6 repress them, suggesting that BR/IAA signalling may determine endocarp cell morphology downstream of the FUL-SHP-AP2 module. By contrast, all three factors repress *SHOU4*, thereby likely promoting secondary cell wall formation in the endocarp. Our data thus show that the FUL-SHP-AP2 module is important in early petunia fruit development to define the endocarp layers, with the morphology of the cells depends on the antagonistic interactions between PhFUL on one hand and ROB1/2/3 and FBP6 on the other hand. Later in fruit development, impaired lignin accumulation and differential regulation of important suberin and lignin biosynthesis genes suggests that the PhFULs, ROB1/2/3 and FBP6 contribute to proper lignification of the endocarp layers. The dehiscence failure in both *fbp6* and *rob1/2/3* fruits illustrates the importance of the *SHP* and *AP2* (co-)orthologs for seed dispersal.

## Materials and Methods

### Plant material & growth conditions

Petunia (*Petunia x hybrida*, line W138) plants were grown in soil (FAVORIT-argile 10) in growth chambers (settings: 16-h day, 22°C/8-h night, 18°C, 75-W Valoya NS12 LED bars; light intensity, 130 mE) at ENS Lyon. The dTPH1 transposon mutants in W138 background (Vandenbussche *et al*., 2003; Vandenbussche *et al*., 2008; Morel *et al*., 2017; Morel *et al*., 2019) were previously described as well as the *pmads3* generated crispr-cas9 alleles (Morel *et al*., 2018). The 35S:FBP6 and 35S:pMADS3 transgenic lines have been generated previously (Heijmans *et al*., 2012a). The genotypes of all single, double, and higher order mutants were confirmed by PCR before anatomical analysis using segregation primers included in Supplementary Table S1.

### Histochemical analysis of fruit sections

WT and mutants were grown next to each other in a growth room to avoid the influence of environmental fluctuations. Flowers of mutant and wild-type plants were hand-pollinated with wild-type pollen and tagged. Fruits were collected at different days after pollination (0, 3, 7, 10, 14, and 21 DAP) and fixed in FAA (3.7% formaldehyde; 5% glacial acetic acid; 50% ethanol) using vacuum-infiltration followed by overnight incubation. For light microscopy and sectioning, the fixed material was dehydrated through a series of ethanol (70%, 95%, 100%, 100%) and further prepared for embedding with an alcohol-toluene series in a Leica automatic tissue processor. The material was then embedded in Paraplast X-tra. The samples were sectioned at 10– 25 μm with an HM 355S rotary microtome (Microm Microtech, France) and stained with safranin/alcian blue, phloroglucinol or toluidine blue. For Johansen’s safranin/alcian blue staining: 2% safranin w/v in 50% ethanol (lignin staining) and 1% Alcian Blue in 3% acetic acid (to observe parenchymatous tissue) was applied (Johansen, 1941). Sections were mounted on Permount and imaged with a Zeiss Axioplan compound microscope equipped with a Axiocam 503 Color Zeiss digital camera with ZEN software. Phloroglucinol staining was performed according to the Cold Spring Harbor Protocol of Sarah Liljegren (http://cshprotocols.cshlp.org/), with the exception that the samples were mounted in a solution of 30% glycerol, 30% HCl, 60% MQ solution to preserve the staining. For Toluidine blue staining, the paraplast embedded sections went through a xylene – ethanol -MQ series (100% xylene for 20’; 100% xylene 2’; 50/50% xylene/ethanol 10’; 100% ethanol 10’, 100% ethanol 2’; 95% ethanol 1’; 95% ethanol 1’; 85% ethanol 1’; 70% ethanol 1’; 50% ethanol 1’; 30% ethanol 1’; 10% ethanol 1’; MQ 1’; MQ 1’) and were then incubated in 0.05% toluidine blue (in MQ) for 10’. After rinsing 3x with MQ, the samples were mounted in 30% glycerol and imaged.

### qRT-PCR analysis

Total RNA was extracted using the Spectrum Plant Total RNA kit (Sigma-Aldrich) and treated with Turbo DNA-free DNase I (Ambion). RNA was reverse transcribed using Maxima Reverse Transcriptase (Thermo Scientific) in presence of RiboLock RNase Inhibitor (Thermo Scientific) according to the manufacturer’s protocol. PCR reactions were performed in an optical 384-well plate in the QuantStudio 6 Flex Real-Time PCR System (Applied Biosystems), using FastStart Universal SYBR Green Master (Roche), in a final volume of 10 µL, according to the manufacturer’s instructions. Primers were designed using the online Universal ProbeLibrary Assay Design Center (Roche) or the online PrimerQuest Tool (IDT). The qRT-PCR profile was as follows: 95°C for 10min, 40 cycles of 95°C for 10s, and 60°C for 30s. Data were analyzed using the QuantStudio 6 and 7 Flex Real-Time PCR System software v1.0 (Applied Biosystems). Petunia *GAPDH* and *RAN* were used as reference genes. PCR efficiency (E) was estimated from the data obtained from standard curve amplification using the equation E = 10−1/slope. Relative expression (R.E.) values on the y axes are the average of three biological replicates normalized to *GAPDH*. All primer sequences are listed in Supplemental Table S1.

### In situ hybridisation of mRNA

Gene-specific sense and antisense RNA probes were synthesised using T7 and SP6 RNA polymerases from PCR-amplified fragments of pGEM-T(Promega)-cloned *PFG, FBP26,* and *FBP29,* 3’ fragments (see Supplemental Table S1). Developing fruits were harvested at 3, 5 or 7 days after pollination (DAP), fixed, and embedded in paraffin using an automated tissue processor (Leica TP1020). *In situ* hybridisation using digoxigenin-labelled probes was performed as described by (Gómez-Mena & Roque, 2018). All antisense probe experiments were repeated at least twice with comparable results. A *MtYUC1* probe was included as negative control in addition to the sense probe to assess background staining.

### Transcriptome profiling of wild-type and *quad* gynoecium and 7 DAP pericarp

Tissue was isolated from gynoecia of 1 cm floral buds and from the pericarp of fruits 7 days after pollination (7 DAP) of WT and *pfg fbp26 fbp29 euap1* (*quad*) fruits. Total RNA was extracted using the Spectrum Plant Total RNA kit (Sigma-Aldrich) and treated with Turbo DNA-free DNase I (Ambion). Library preparation and Illumina sequencing were performed at BGI Genomics (stranded library; Illumina NovaSeq 2×150 nt Paired End sequencing). After quality control, the filtered reads were mapped to the *Petunia axillaris* genome sequence v1.6.2 (Bombarely *et al*., 2016) and all data were analyzed using the CLC work package using default settings. In the PCA analysis (Metsalu & Vilo, 2015), we noticed that *quad* sample two behaved like the WT at the PC1 axis (Supplementary Fig. S8). To find an explanation for this, we performed a differential expression analysis and scanned by eye the RPKM values to find out where q_7_2 was similar to the WT 7 DAP samples. This revealed that many circadian rhythm associated genes, such as homologs of the clock genes *LHY* (Peaxi162Scf00092g01520) and *LCL5* (Peaxi162Scf00472g00610) were differentially regulated for the WT samples plus q_7_2 versus the other two *quad* samples. Thus, PC1 reflected diurnal expression differences rather than differences between WT and *quad* (due to the irregular sampling of the *quad* samples, where time of the day had not been sufficiently taken into account). Of the original list of 1972 DEGs (FDR p-value <0.01), 242 genes displayed an expression value for q_7_2 that was more similar to the WT samples than to q_7_1 and q_7_3. These genes were omitted from the DEG list. For the gynoecium samples, we did not identify a clock-related bias. For data validation with qRT-PCR, new batches of plants were grown and pericarp tissues were isolated from fruits at 0 DAP, 7 DAP and 14 DAP from WT and different mutant backgrounds. For each biological replicate, tissues harvested from three fruits originating from three independent plants of the same genotype were pooled. RNA extraction, cDNA synthesis and qPCR were performed as described above (see Supplemental Table S1 for the primers). The raw data from the RNA-Seq experiment has been uploaded to the European Nucleotide Archive (ENA, https://ena-docs.readthedocs.io/en/latest/retrieval/general-guide.html) under project PRJEB99017 (Note: the gynoecium samples are referred to as ‘0 DAP’ in this submission).

### Electrophoretic Mobility Shift Assay

*PFG* and *FBP23* coding sequences were derived from previously generated pDEST32 vectors (Immink *et al*., 2003; Ferrario, 2004) and recombined in the pSPUTK vector using the Gateway system (Invitrogen). The pSPUTK vector allows *in vitro* protein synthesis using the TnT SP6 High-Yield Wheat Germ Protein Expression System (Promega) according to the manufacturer’s instructions. The probe fragments were amplified from petunia genomic DNA (80–120 bp containing the in silico predicted CArG-boxes approximately in the middle, see Fig. 3D for the probe locations) and cloned into the pJET vector. EMSAs were performed as described (Smaczniak, C. *et al*., 2012). Labelled probe fragments were obtained by PCR using DY-682 primers that flank the fragment inserts in the pJET backbone. Labelled fragments were then purified with the NucleoSpin Gel and PCR Clean-up kit (MACHEREY-NAGEL). Gel shifts were visualized using a LiCor Odyssey imaging system at 700 nm.

## Results

### Petunia *AP1/FUL* genes redundantly determine carpel number and placenta size

The Arabidopsis genome encodes four members of the AP1/FUL clade, but only one of them, FUL, plays a major role in fruit development (Gu *et al*., 1998a). In an earlier study (Morel et al, 2019), we found that the four petunia *AP1/FUL*-like genes function in a largely redundant fashion in inflorescence and flower development. We therefore hypothesized that their functional overlap may extend to fruit development. As a first test, we monitored their expression levels in fruits at several stages of development using qRT-PCR (Fig. 1A). Fruits from 7 DAP (days after pollination) onwards were dissected into pericarp and remaining tissues (placenta + ovules). Overall, this revealed that all four genes continue to be expressed throughout fruit development, although *euAP1* at a considerably lower level than the *FUL*-like genes. The three *FUL*-like genes have comparable expression levels and are each much higher expressed in the pericarp than in the placental + ovule tissues. To reveal a possible role in fruit development, we examined fruit morphology of 14 DAP fruits obtained from *AP1/FUL* double mutants (*pfg fbp26*, *fbp29 fbp26*, *fbp26* e*uap1*, *pfg euap1 and fbp29 euap1* (Morel *et al*., 2019)). This did not yield any obvious aberrant fruit phenotype, except for moderate variation in fruit sizes, while pericarp sections appeared normal (Supplementary Fig. S1). This indicates that the petunia *AP1*/*FUL*-like genes may act redundantly during fruit development similar to their functions earlier in reproductive development. We then investigated the fruits of the higher order mutants *euap1 fbp26 pfg*; *euap1 fbp26 fbp29*, *fbp26 fbp29 pfg*; *euap1 fbp29 pfg*, and the quadruple mutant *euap1 fbp26 fbp29 pfg* (hereafter called ‘*quad*’). Interestingly, *quad* mutants had a prominent fruit phenotype clearly visible by eye: while overall maturation and dehiscence of the *quad* mutant fruits appeared normal, with dehiscence occurring around 4-5 weeks after pollination, its fruits were consistently larger than wild type (Fig. 1B-C). We observed that the majority of the *quad* mutant fruits were composed of three fused carpels or more, instead of two carpels normally found in wild type (Fig. 1B, 1D). The number of floral organs, including the number of carpels, is established very early during flower development, when the floral organ primordia are specified from the floral meristem, and we therefore expect that this function of the *AP1/FUL* genes is rather linked to their developmental role preceding fruit initiation. Indeed, Morel et al. (2019) described that *quad* mutant flowers have typically more floral organs. Here, we quantified this and found that floral organ number was increased in all four whorls of the *quad* mutant, with 28 out of 36 flowers analysed exhibiting three carpels or more, while the wild type consistently has two (Fig. 1D). We observed this high frequency of increased carpel number only in *quad* mutants, indicating that the four petunia *AP1/FUL-like* genes redundantly are required to assure fixed floral organ numbers. While this increase in carpel number may contribute to the size increase of the *quad* fruits, we found that *quad* fruits with two carpels were also larger due to a major increase in placenta size (Supplementary Fig. S2). In addition, this major placenta size increase was also observed in *euap1 fbp26 fbp29* fruits that are typically composed of two carpels (Fig. 1C, Supplementary Fig S2). We measured fruit diameter for the different triple mutant combinations and found a significant size increase for *euap1 fbp26 fbp29* fruits, comparable to the *quad* fruits. For the other triple combinations, large within-genotype variation was observed, indicating that placenta size is less robustly regulated in those lines (Fig. 1C). Together, our data show that the observed fruit size increase is mainly due to a larger placenta, with euAP1, FBP26 and FBP29 redundantly limiting its size in wild type.

**Figure 1.**
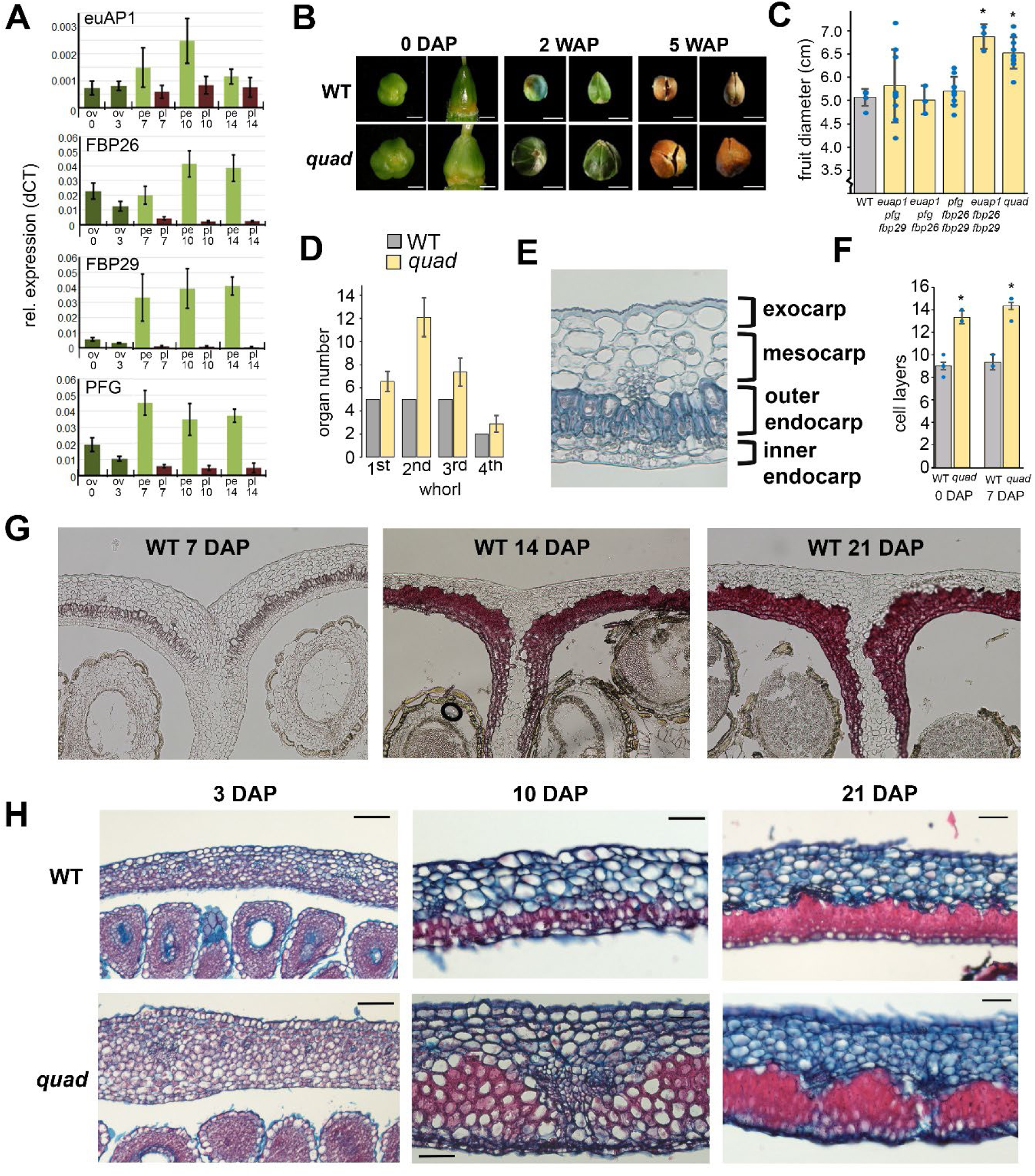
Characterization of *quad* mutant fruits. A) Relative expression profiles of the petunia AP1/FUL-like genes in fruits obtained by qRT-PCR. The numbers below the bars indicate the number of days after pollination. The letter abbreviations indicate the tissue type: ov=entire ovary, pe=pericarp, pl=placenta + ovules. The error bars indicate the SE based on 3 biological replicates. B) Macroscopic phenotype of WT and *quad* mutant fruits at 0 days after pollination (0 DAP=unpollinated), 2 weeks after pollination (2 WAP), and 5 WAP. Scale bars: 0 DAP=1mm; 2/5 WAP=5mm. C) Quantification of fruit diameter in different mutant combinations. The blue dots represent the individual data points. Significance is indicated with an asterisk (T-test, p<0.05). D) Quantification of floral organ number in the different whorls (1^st^, sepals, 2^nd^, petals; 3^rd^, stamens; 4^th^, carpel). The error bars indicate the standard deviation based on 10 flowers (WT) or 36 flowers (*quad*). E) Section of WT pericarp at 7 DAP, stained with toluidine blue. The different tissue types are indicated. F) Quantification of cell layer number in pericarp sections of WT and quad mutants at 0 and 7 DAP. Significance is indicated with an asterisk (T-test, p<0.05). G) Sections of WT pericarp at 7, 14 and 21 DAP at the junction of the two carpels. Sections are stained with phloroglucinol, which visualizes lignification in dark red. H) Safranin/alcian blue-stained sections of WT and *quad* fruits of different stages. Scale bars: 50 *μ*m.

### The petunia *AP1*/*FUL*-like genes additively regulate pericarp development

To get more insight in the anatomy of the *quad* mutant fruits, we sectioned fruits at different stages of development and compared with the wild type. As described by

Pabon-Mora and Litt (2011), the capsular petunia fruit consists of two fused carpels, and the inner placenta connects to the carpels at the fusion sites (sometimes called septum). At this location, small cell files develop during fruit development to allow dehiscence of the upper part of the mature fruit. The pericarp contains four different types of cell layers, which are (from outer to inner): the single-layered exocarp, the mesocarp layers containing vascular bundles, the outer endocarp layers and the inner endocarp layers (Fig. 1E). Dependent on the position in the fruit (apex, mid or base), the number of layers from each type and the degree of lignification in the endocarp layers varies in wild-type fruits (Pabón-Mora & Litt, 2011). Therefore, to avoid these variations, we systematically sectioned the mid-region of fruits, unless mentioned otherwise. Wild-type W138 fruits (the genetic background of the transposon lines) contain a single layer of exocarp cells at seven days after pollination (7 DAP), 4-6 layers of roundish parenchymatous mesocarp cells, two layers of narrow sclerenchyma cells that form the outer endocarp, and two layers of smaller, more rectangular cells that form the inner endocarp (Fig. 1E-F). Phloroglucinol staining revealed that in the mid region, the outer endocarp layers of the wild-type fruits begin to lignify around 7 DAP, which is fully completed at 14 DAP (Fig. 1G). The inner endocarp layers develop thicker walls and initiate lignification only around 14 DAP, which increases until 21 DAP (Fig. 1G). At the carpel fusion zones, the endocarp layers bend inwards to connect with the placenta, leaving several cell files unlignified, which mark the future dehiscence zone. To thoroughly monitor differences in cell layer structure, cell morphology, and lignification between the pericarp of wild-type and *quad* fruits, we used safranin/alcian blue staining and sectioned fruits at 0, 3, 7, 10, 14 and 21 DAP (Figure 1H and Supplementary Fig. S3). The sectioning revealed that in addition to having a larger placenta, *quad* mutant fruits also contain more pericarp layers. At 0 and 7 DAP, the wild-type ovary wall consists of 9 to 10 cell layers, whereas the ovary wall of *quad* mutants consists of 13 to 14 cell layers, with the number of cell layers in both genotypes being only slightly higher at 7 DAP compared to 0 DAP (Fig. 1F, H).). Lignification starts in both WT and *quad* fruits around 7 DAP in the outer endocarp layer, and reaches the inner endocarp layers between 10 and 14 DAP in the wild type. However, in *quad* fruits, lignification in the inner endocarp layers hardly occurs and the lignification in the outer endocarp layers is patchy (Fig. 1G-H, Supplementary Fig. S3). Furthermore, both cell arrangement and cell shape in the inner and outer endocarp layers differ from the wild type. Whereas the wild-type endocarp cells have a regular arrangement and are clearly smaller than the mesocarp cells, the *quad* cells are chaotically arranged and are rounder and larger than the wild type, more reminiscent of mesocarp cells. This suggests that the petunia *AP1/FUL*-like genes are required for proper endocarp development.

To obtain insight in the contributions of the different *AP1/FUL-*like genes to the observed fruit phenotypes in the *quad* mutant, we inspected in detail the pericarp phenotypes of the four triple mutant combinations, using sections of fruits from 0, 3, 7, 14 and 21 DAP stained with toluidine blue, safranin/alcian blue or phloroglucinol. Due to a lower number of flowers in higher-order *PhFUL* mutants (Morel et al., 2019), not all stages were sampled for all triple mutants. Interestingly, the *pfg fbp26 fbp29* mutant has the most severe pericarp phenotype (Fig. 2A-D). In addition to containing more pericarp cell layers than the wild type (Fig. 2E), endocarp cells are always much larger and rounder and closely resemble the phenotype of the *quad* mutant. Thus the establishment of endocarp cell morphology, which occurs around 3-5 DAP, is mainly regulated by the three *FUL*-like genes, while *euAP1* plays at most a minor role. In agreement with the phenotype of the *quad* mutant, the endocarp of *pfg fbp26 fbp29* mutants was less lignified at 7 DAP, and lignification was patchy at 14 DAP (Figs. 2B-2D). Other triple combinations displayed somewhat milder pericarp phenotypes and we observed substantial variation between fruits of the same genotype (Fig. 2A-D, Supplementary Figs. 4-5), again suggesting that reduced dosage of the *AP1/FUL*-like genes results in an imbalance with a variable phenotypic effect. In conclusion, our data indicate that *PFG*, *FBP26,* and *FBP29* function redundantly in the regulation of pericarp patterning and endocarp development, in line with their higher expression level compared to *euAP1* (Fig. 1A). To get more insight in the spatial transcript accumulation of *PFG*, *FBP26* and *FBP29* within the pericarp, we attempted to perform in situ hybridisations with embedded tissues of 7 DAP wild-type fruits. However, although a very clear signal appeared in the endocarp layers, this signal was equally clear in the samples that were treated with the sense probe. This was visible already prior to lignification of the endocarp layers, suggesting a cross-reaction between lignin precursors and the alkaline phosphatase reporter enzyme in the assay (Supplementary Fig. S6). In a second attempt, we performed the *in situ* hybridisation for *PFG* and *FBP26* with 3 and 5 DAP samples. In the sections from these stages, the background signal was absent in the inner endocarp, while the signal in the outer endocarp depended on stage.At 5 DAP, background was clearly apparent, while this was not the case at 3 DAP (Fig. 2F). However, signals were strong in the outer endocarp layer when the *PFG* and *FBP26* antisense probes were used (Fig. 2F), suggesting that both genes are distinctly expressed in this layer, in agreement with the observed mutant phenotypes. Furthermore, signal was observed in the inner endocarp layer, particularly for *PFG*, but not in the mesocarp layer (Fig. 2F). This indicates that the *FUL*-like genes are predominantly expressed in the endocarp layers of the petunia fruit. Thus, the combination of our expression and phenotypic analyses suggest that *PFG*, *FBP26* and *FBP29* are functioning in the endocarp layers to regulate their development at an early stage and contribute to the regulation of secondary metabolite accumulation (e.g. lignin) at a later stage.

**Figure 2.**
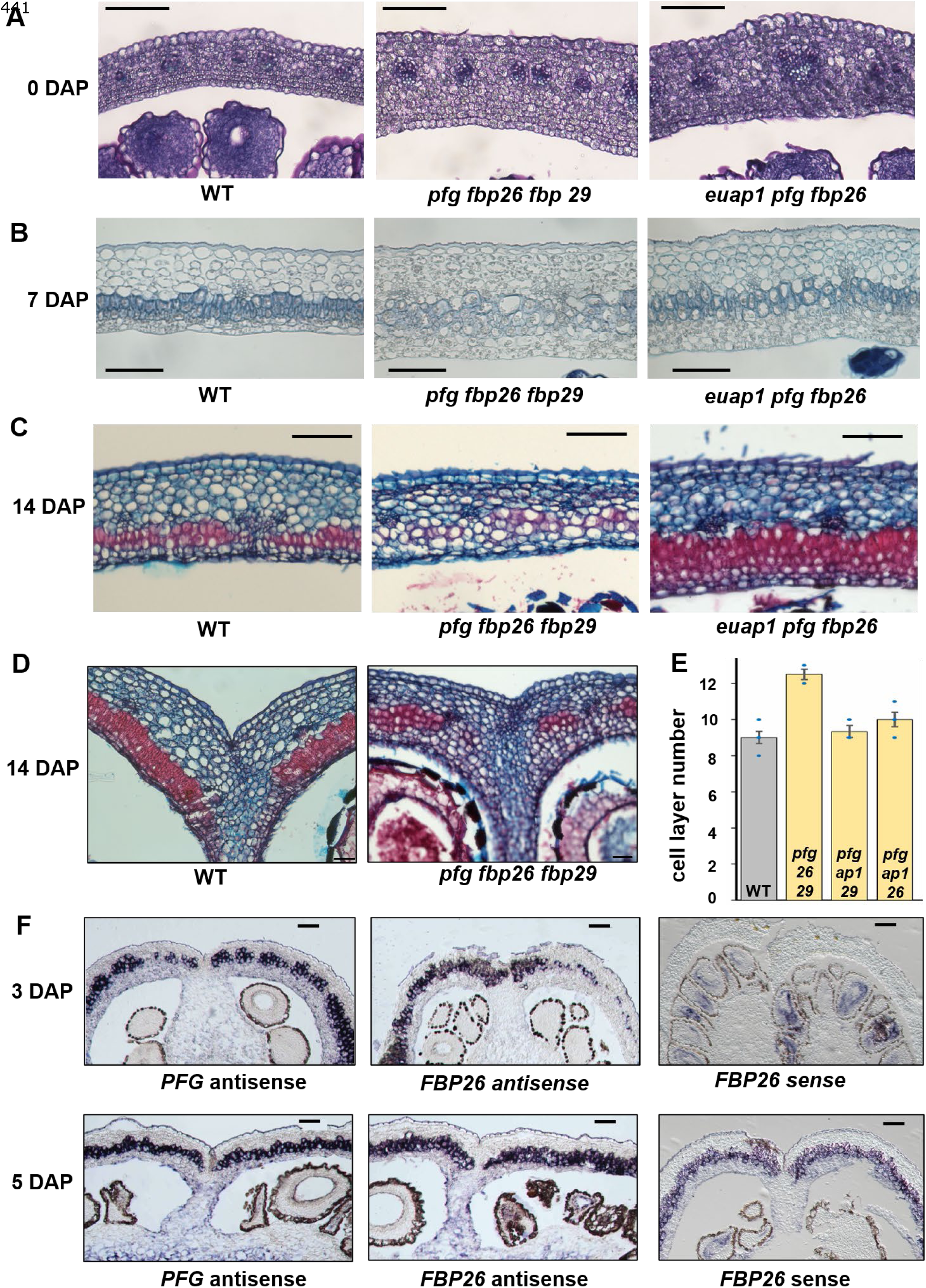
**Contributions of the four petunia *FUL/AP1* genes to pericarp patterning**. A-C) Sections of pericarp at 0 DAP (unpollinated) (A), 7 DAP (B) and 14 DAP (C), stained with toluidine blue (A-B) or safranin/alcian blue (C). The genotypes are indicted below the picture frames. Additional pictures are available in Suppl. Figures S4-S5. D) Safranin/alcian blue-stained sections of 14 DAP fruits at the junction of the two carpels. E) Quantification of cell layer number in pericarp sections of WT and triple mutants at 0 DAP. F) Pictures of an *in-situ* hybridization experiment with 3 and 5 DAP pericarp sections. Upper panels: 3 DAP sections hybridized with PFG antisense (left), FBP26 antisense (middle) and FBP26 sense probes. Lower panels: 5 DAP sections hybridized with the same probes. Additional pictures, including the PFG sense probe, are available in Suppl. Fig. S6. Scale bars A-C, F: 100 *μ*m. Scale bars D-E: 50 *μ*m.

**Figure 3.**
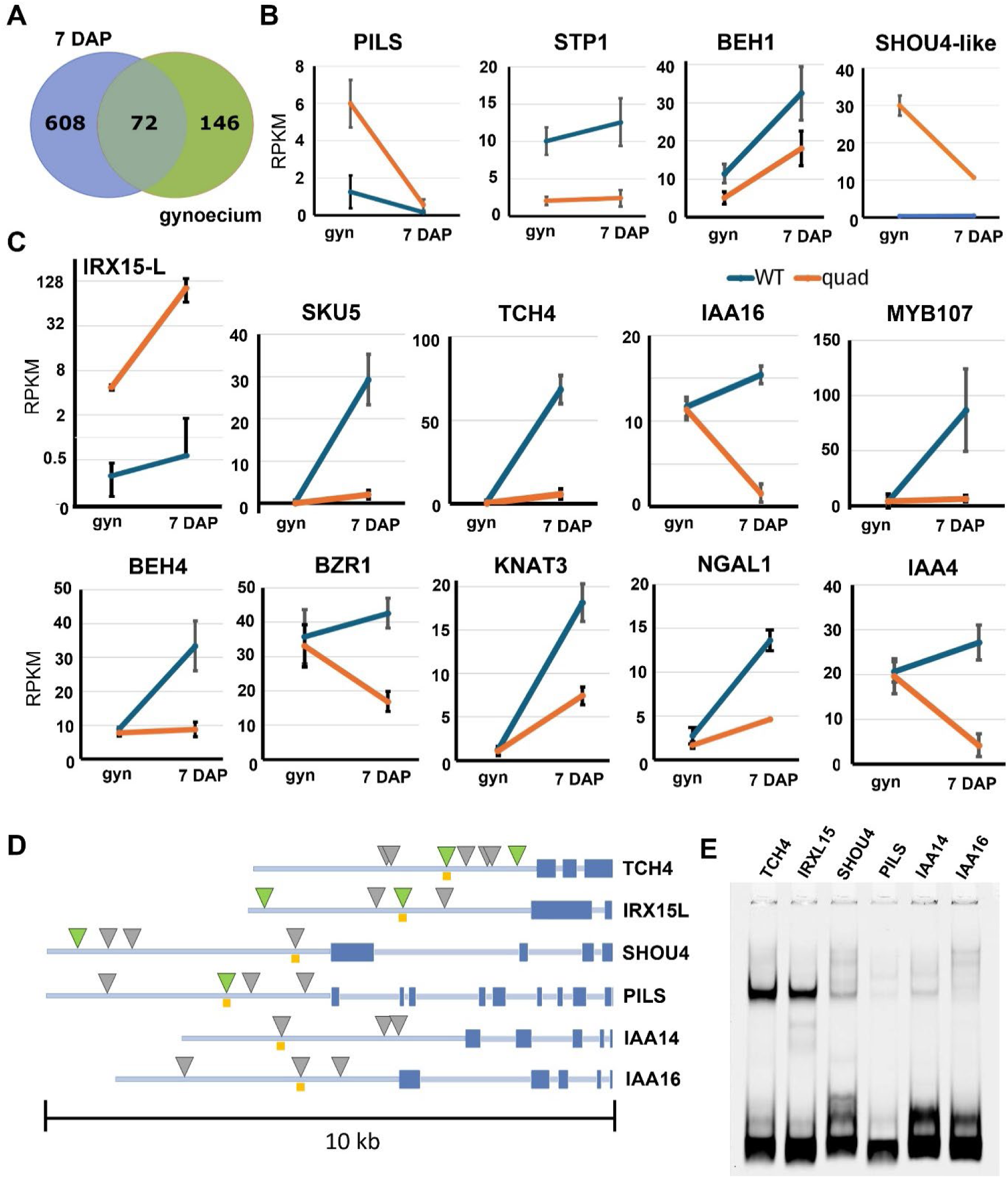
**Downstream targets of the *PhFUL* genes in the pericarp**. A) Venn diagram showing the verlap in the DEGs from gynoecium (WT vs *quad*) and 7 DAP (WT vs *quad*). B) RPKM plots based on he RNAseq results from a selection of interesting targets from the gynoecium list. Both the expression in he gynoecium and the 7 DAP fruit is depicted. Blue lines indicate the wild-type levels, orange lines the *uad* levels. C) RPKM plots based on the RNAseq results from a selection of interesting targets from the DAP list. Blue lines indicate the wild-type levels, orange lines the *quad* levels. D) (next page) D) Gene structures of genes tested in (E). The narrow faint blue bar indicates the promoter region and introns; the thicker blue bars the exons. CArG-box-like sequences are indicated with grey triangles. Canonical CArG-boxes, generally bound with high affinity (Aerts et al, 2018; van Mourik et al, 2023; Thoris et al. 2025), are indicated with light green triangles. The orange boxes indicate the probe fragments used in (E). The black bar below indicates the scale. E) Electrophoretic Mobility Shift Assay (EMSA) of DNA probe fragments of 6 different target genes tested with in-vitro produced PFG-FBP23 complexes. The black bands at the bottom indicate the non-bound DNA probes; bands higher in the gel (clearly visible for the TCH4 and IRXL15 probes) indicate shifted probes due to binding of the PFG-FBP23 complex.

### Identification of downstream genes associated with the *quad* mutant phenotypes

To further investigate how the petunia *FUL*-like genes perform their functions during fruit development, we performed an RNA-seq experiment in which the transcriptome of *quad* mutant tissue was compared with that of the wild type. We selected two different tissue types: i) gynoecium of 1cm floral buds, to capture possible transcriptomic changes during gynoecium development that may underlie early-established phenotypes such as ovary wall layer number, and ii) pericarp of 7 DAP fruits to identify differentially expressed genes involved in pericarp patterning and endocarp cell functioning. High-throughput sequencing yielded 32-36 million reads per sample, and a principal component analysis was performed to evaluate the clustering of the samples. While the WT samples showed clear clustering, the *quad* samples were more dispersed (Supplementary Fig. S7). Reason for this is probably the scattered sampling due to the very low production of flowers in the *quad* mutant. Analysis revealed that there was also a circadian bias, particularly in the 7 DAP samples, and we corrected for this (see Materials & Methods). A list of 1732 DEGs at FDR <0.01 or a more stringent set of 680 at Bonferroni p-value <0.05 was obtained (Supplementary Table S2). A much smaller list of 457 (FDR p-value < 0.01) and 218 (Bonferroni p<0.05) DEGs was identified for the gynoecium samples, consistent with the minor differences between WT and *quad* fruits at this early stage (Supplementary Table S3). 72 out of these 218 DEGs were overlapping with the 7 DAP DEG set (Fig. 3A; Supplementary Table S4), and of the overlapping genes, 49% was downregulated in the *quad* mutant, while 51% was upregulated.

To decipher the function of the *PhFUL* genes in pericarp development further, we examined the annotations of the identified DEGs. A GO-term enrichment analysis revealed several enriched ‘Biological function’ categories for the 7 DAP samples, among which many associated with cell growth and development, as well as with cell-wall synthesis and biosynthetic processes involved therein (Supplementary Table S5a). These include ‘regulation of cell morphogenesis’ (GO:0022604), ‘pectin catabolic process’ (GO:0045490), ‘galacturonan metabolic process’ (GO:0010393), and ‘cell wall organization or biogenesis’ (GO:0071554), but also the categories ‘regulation of DNA-templated transcription’ and ‘response to hormones’ were enriched. The list of enriched categories was much shorter for the gynoecium samples, and did not include any category that seemed associated with the observed phenotypes (Supplementary Table S5b).

Therefore, we carefully inspected the list of gynoecium DEGs for interesting target genes with a considerable change in expression. We identified several genes with annotations that may be linked to the increased number of pericarp layers (Fig. 3B, Supplementary Fig. S8, Supplementary Table S6). This includes an auxin transporter that belongs to the PIN-LIKES (PILS) (Peaxi162Scf00385g00415). PILS proteins act in the endoplasmic reticulum to repress transport of auxin into the nucleus, regulating intracellular auxin homeostasis (Barbez *et al*., 2012; Sun *et al*., 2020) and can thus render cells more or less sensitive to auxin. Auxin signalling in the *quad* gynoecium may also be altered by the downregulation of a close homolog of *IAA14* (Peaxi162Scf01416g00118) and upregulation of *ARF11* (Peaxi162Scf00747g00018) (Supplementary Table S6). In addition to modified auxin signalling, brassinosteroid (BR) signalling appears different as well, as a close homolog of *BES1/BZR1*-homolog *1* (*BEH1,* Peaxi162Scf00064g00329), which regulates the brassinosteroid response in Arabidopsis redundantly with five other BZR1-like factors (Chen *et al*., 2019), is downregulated. Another interesting DEG that may be associated with cell division/patterning, is Peaxi162Scf01019g00112, a homolog of *Arabidopsis Sugar Transporter 1* (*STP1*), which regulates the uptake of monosaccharide sugars (Flütsch *et al*., 2020). Possibly, the crosstalk between sugar and auxin, with glucose for example being required for correct PIN2 localisation (Mishra *et al*., 2022), plays a role in the observed phenotype. Finally, Peaxi162Scf00442g00524, a homolog of Arabidopsis *SHOU4*-like, is highly upregulated in the *quad* gynoecium (Fig. 3B). SHOU4-like proteins negatively regulate cellulose synthases (CESAs) by controlling their trafficking at the plasma membrane. As such, they influence the direction and extent of cell wall extension and thereby probably cell division (Polko *et al*., 2018). Thus, our data suggest that the *AP1/FUL*-like genes influence auxin, BR and sugar signalling, as well as regulate cellulose deposition, possibly to control periclinal cell divisions in the ovary wall and/or to regulate placenta size.

### The PhFULs regulate secondary cell wall formation at 7 DAP

While the gynoecium DEG list contained relatively few interesting genes, a close inspection of the 7 DAP DEG list revealed many deregulated genes with relevant annotations, in line with the GO-term analysis. These genes could be grouped into several classes (Supplementary Table S6): I) (secondary) cell wall formation, II) regulation of lignin biosynthesis, III) hormone signalling, and IV) transcription factors (TFs). Because the phenotype of the *quad* fruits at 7 DAP is markedly different from that of the wild type, many DEGs may reflect the endocarp development defect, rather than being direct targets. This is probably coinciding with changes in secondary cell wall formation and lignification, common features of the endocarp in dry dehiscent fruits (Dardick & Callahan, 2014). Amongst the genes associated with secondary cell-wall formation, the strong upregulation of a close homolog of IRREGULAR XYLEM 15-like (IRX15-L) (Peaxi162Scf00549g00028), responsible for deposition of the hemicellulose xylan in the secondary cell wall (Brown *et al*., 2011), is most remarkable. In the wild-type pericarp, the expression of this gene is significantly induced at the 7 DAP stage but its induction in the *quad* mutant is enormous (Fig. 3C). Thus, it appears that the PhFULs repress *IRX15-L* to temper the deposition of xylan in the secondary cell wall at 7 DAP. The *quad* endocarp cells are more chaotically arranged than the WT, are larger and have irregular cell shapes, reminiscent of mesocarp cells. This phenotype may be explained by the downregulation of a close homolog of the multicopper oxidase-like protein skewed 5 (SKU5), Peaxi162Scf01181g00022. Arabidopsis SKU5 and its family members, the SKU5-like (SKS) proteins, are GPI-anchored proteins that link cell wall synthesis to cell expansion (Quinn et al., 2023; Chen et al., 2024). Mutants exhibit a large variation in cell sizes, as well as aberrant cell walls and division planes (Zhou, 2019; Chen *et al*., 2023; Quinn *et al*., 2024). Petunia *SKU5-like* is strongly induced in the wild-type 7 DAP pericarp, in agreement with the timing of endocarp differentiation, but not at all in the *quad* pericarp (Fig. 3C). A similar expression pattern is detected for *TOUCH4*-like (*TCH4*-like, Peaxi162Scf00111g00022, Fig. 3C), which encodes in Arabidopsis a cell-wall modifying enzyme that regulates pectin deposition in the primary cell wall (Zhang *et al*., 2022). The combination of promoting cell-wall organisation via *SKU5*-like and *TCH4*-like, but repressing secondary cell wall deposition via *IRX15*-like, suggests that the PhFULs regulate endocarp differentiation at 7 DAP, while preventing rapid secondary cell wall formation in these cells. This may particularly be relevant in the inner endocarp cells, which lignify only at a late stage.

Given the putative dual role of the PhFULs in the promotion of secondary cell wall formation, but retardation of secondary cell metabolite deposition, we next looked at genes involved in lignin deposition. In the *quad* pericarp, lignin deposition is patchy in the outer endocarp cells (Figs. 1G), and reduced in the inner endocarp layers at 21 DAP (Supplementary Fig. S3), suggesting that the PhFULs directly or indirectly regulate lignin deposition. Notably, a set of MYB transcription factors that is associated with phenylpropanoid (a lignin precursor) and lignin biosynthesis, is differentially regulated in the *quad* pericarp. These include both putative activators (close homologs of Arabidopsis *MYB15*, *MYB42*, *MYB43* and *MYB58*) and a putative repressor (MYB3 homolog) of phenylpropanoid/lignin biosynthesis (Supplementary Table S6, Supplementary Fig. S8) (Zhou *et al*., 2009; Chezem *et al*., 2017; Zhou *et al*., 2017; Geng *et al*., 2020; Choi *et al*., 2023). Therefore, the effect of the differential regulation of these *MYBs* on endocarp lignification, and the role of the PhFULs therein is not yet clear. Possibly, the PhFULs also balance lignin deposition via the *MYBs*. We additionally checked upstream regulators of secondary cell wall development (close homologs of *NST1/2* and *VND6/7*), but did not find significant differences. Interestingly, also the putative ortholog of *MYB107* (Peaxi162Scf00006g00098), an important upstream regulator of the suberin pathway, is strongly downregulated in the *quad* mutant (Fig. 3C). In the wild-type pericarp, this gene is significantly induced at the 7 DAP stage, but this induction is almost absent in *quad*, indicating that the PhFULs also strongly promote, directly or indirectly, the biosynthesis of suberin, a cell-wall modifier that provides a barrier to prevent water loss and pathogen ingrowth (Woolfson *et al*., 2022).

The third class of DEGs clearly deregulated in the *quad* pericarp contains hormone-signalling genes. Four AUX/IAA genes, homologs of *IAA4*, *IAA14* and *IAA16*, which inhibit auxin response factors, are considerably expressed in the wild-type pericarp but have much lower expression levels in the *quad* pericarp (Fig. 3C, Supplementary Fig. S8). Furthermore, the extracellular auxin transporters Peaxi162Scf00683g00461 and Peaxi162Scf00253g01129 (close homologs of *PIN1* and *PIN3*, respectively) are highly expressed in the gynoecium of both wild type and *quad*, but strongly downregulated at 7 DAP (Supplemental Fig. S8), with significantly lower levels in *quad*. Next to auxin signalling, also brassinosteroid (BR) signalling is altered in the *quad* mutant at 7 DAP, as in the gynoecium stage. However, instead of *BEH1*, whose differential expression is just not significant at this stage, close homologs of *BEH4* (Peaxi162Scf00112g01217) and *BZR1* (Peaxi162Scf01327g00017) are upregulated (Shi *et al*., 2022). While both genes are strongly induced in the wild-type pericarp at 7 DAP, this induction is completely absent in the *quad* mutant (Fig. 3C). Taken together, our transcriptome analysis provides a strong indication that both auxin and brassinosteroid signalling are important factors in early pericarp development, and that this signalling is regulated by the PhFULs.

### Several transcription factors act downstream of the PhFULs

Evaluating the functions of the Arabidopsis orthologs/close homologs of the up-and downregulated TF-encoding genes, we identified five bHLH family members, three SPB-box TFs, a class II KNOX homeodomain TF, an AP2/ERF, and an AP2/B3 TF that appeared interesting due to their putative developmental functions (Supplementary Table S6). In particular the differential regulation of the class II KNOX gene Peaxi162Scf01184g00135, closely related to Arabidopsis *KNAT3*, is very interesting. Arabidopsis KNAT3 is a regulator of secondary cell wall formation, which can probably physically interact with the key regulators NST1/NST2 to influence lignin biosynthesis (Qin *et al*., 2020). *knat3 knat7* mutants display an irregular xylem phenotype with reduced xylan content (Wang *et al*., 2020; Nookaraju *et al*., 2022). Petunia *KNAT3* is strongly induced in wild-type pericarp at 7 DAP, but this induction is much weaker in the *quad* pericarp (Fig. 3C). Also notable is the downregulation of the NGATHA-type AP2/B3 TF Peaxi162Scf00096g00099, the putative ortholog of Arabidopsis *NGATHA-like 1* (*NGAL1*). *NGAL1-3* are members of the RAV subfamily, which are controlling organ boundary formation by repressing the activity of the *CUP-SHAPED COTYLEDON* (*CUC*) genes in Arabidopsis (Nicolas *et al*., 2022), a function that may also be needed during pericarp layer differentiation in the petunia fruit (Fig. 3C).

In species such as Arabidopsis and tomato, FUL and its co-orthologs mainly act as repressors (Bemer *et al*., 2017; Jiang *et al*., 2021). Here, we found both strongly upregulated and downregulated genes in the *quad* mutant. To identify loci that may be directly regulated by the PhFULs, we focused on the interesting genes that were strongly upregulated in *quad* mutants. This included the close homologs of *PILS, STP1, SHOU4-like, IAA14/16, IRX15-like, SKU5, TCH4, MYB107, BEH4, BZR1, KNAT3, AGL24,* and *FLC*, for which we analysed the 5 kb upstream region *in silico*. With the exception of *BEH4*, all genes had CArG box-like sequences in their upstream regions. CArG boxes are known to be bound by MADS-domain TFs, with slight differences in the nucleotide sequence of the motif contributing to complex-specific binding (van Mourik *et al*., 2023; Thoris *et al*., 2024). Moreover, *PILS*, *SHOU4*-like, *IRX15*-like, *SKU5* and *TCH4* contain canonical CArG boxes in their upstream region, which included an A/T extension defined by CC[A/T]_6_GG[A/T]_3_, reported to be strongly bound by FUL-AG-SEP3 complexes in the Arabidopsis silique (Aerts *et al*., 2018; Thoris *et al*., 2024). To determine to what extent the PhFULs can bind to the different identified CArG boxes, we tested a subset of upstream fragments using an electrophoretic mobility shift assay (EMSA) (Figs. 3D and 3E). Because PhFUL proteins need to heterodimerize to bind to the DNA, we selected FBP23, a SEPALLATA-like MADS-box protein, as heterodimerization partner expressed in the fruit (Immink *et al*., 2003; Ferrario, 2004). The PFG-FBP23 complex could strongly bind to the canonical CArG boxes present in the promoters of the *TCH4* and *IRX15L* homologs, and weakly to *SHOU4*-like. However, no clear shift was observed for probe fragments of the *PILS*, *IAA14* and *IAA16* homologs, indicating that these are not direct targets of PhFUL, or that PFG binds to regulatory regions not tested in the EMSA. Thus, it is plausible that the PhFULs directly regulate (secondary) cell wall biosynthesis and thereby control endocarp layer differentiation and development, while their effect on hormone signalling may be indirect.

### The *SHP*-ortholog *FBP6* regulates different aspects of fruit development

*FUL*, *SHP/AG* and *AP2* genes regulate aspects of fruit development in both tomato and Arabidopsis (Gu *et al*., 1998b; Ferrándiz *et al*., 2000; Liljegren *et al*., 2000; Vrebalov *et al*., 2009; Chung *et al*., 2010; Karlova *et al*., 2011; Ripoll *et al*., 2011; Bemer *et al*., 2012; Gimenez *et al*., 2016), and we therefore investigated whether a FUL-AP2-SHP module may be conserved in petunia as well. First, we investigated the role of the two C-class genes *(AG/SHP* orthologs*) pMADS3 and FBP6*. We used *dTph1* transposon mutants and crispr-cas9 alleles previously generated (Morel *et al*., 2018). Additionally, we included available *35S:pMADS3* and *35S:FBP6* overexpression lines (Heijmans *et al*., 2012a; Morel *et al*., 2018). Analysing *fbp6 pmads3* double mutants was not possible, because carpels are not formed in this complete C-class knock out (Heijmans *et al*., 2012a). While fruits from *pmads3* mutants appeared completely normal throughout development, resulting in fully dehiscent fruits at maturity, *fbp6* fruits failed to open properly (Fig. 4A-B), even after extended drying of the seedpods. The pods of petunia fruits typically open at the top to shatter the seeds, linked to detachment of the two valves at the location of a few cell files with smaller cells (Fig. 1F). During WT early fruit development, the style, which has a very small diameter compared to the ovary, abscises from the apical fruit region around 3-4 DAP together with petal and stamen abscission. We observed that in *fbp6* mutant fruits, style abscission usually does not occur, (Fig. 4A-B). Occasionally, style abscission occurs late but always well above the normal style abscission point. Earlier, it was already observed macroscopically that *fbp6* pistils develop a much less clear transition zone between ovary and style (Heijmans *et al*., 2012a; Morel *et al*., 2018). This seems to be maintained after fertilization and further fruit development (Fig. 4B). We observed in sections of the apical region of *fbp6* fruits that there is a broader connection between placenta and pericarp than in the wild type, and the cell files with small cells appear absent or much reduced (Fig. 4C). This indicates that *FBP6* is specifically required for the correct patterning of the dehiscence zone in the apical region of the fruit, which may cause its failure to abscise the style as well as its dehiscence defect.

**Figure 4.**
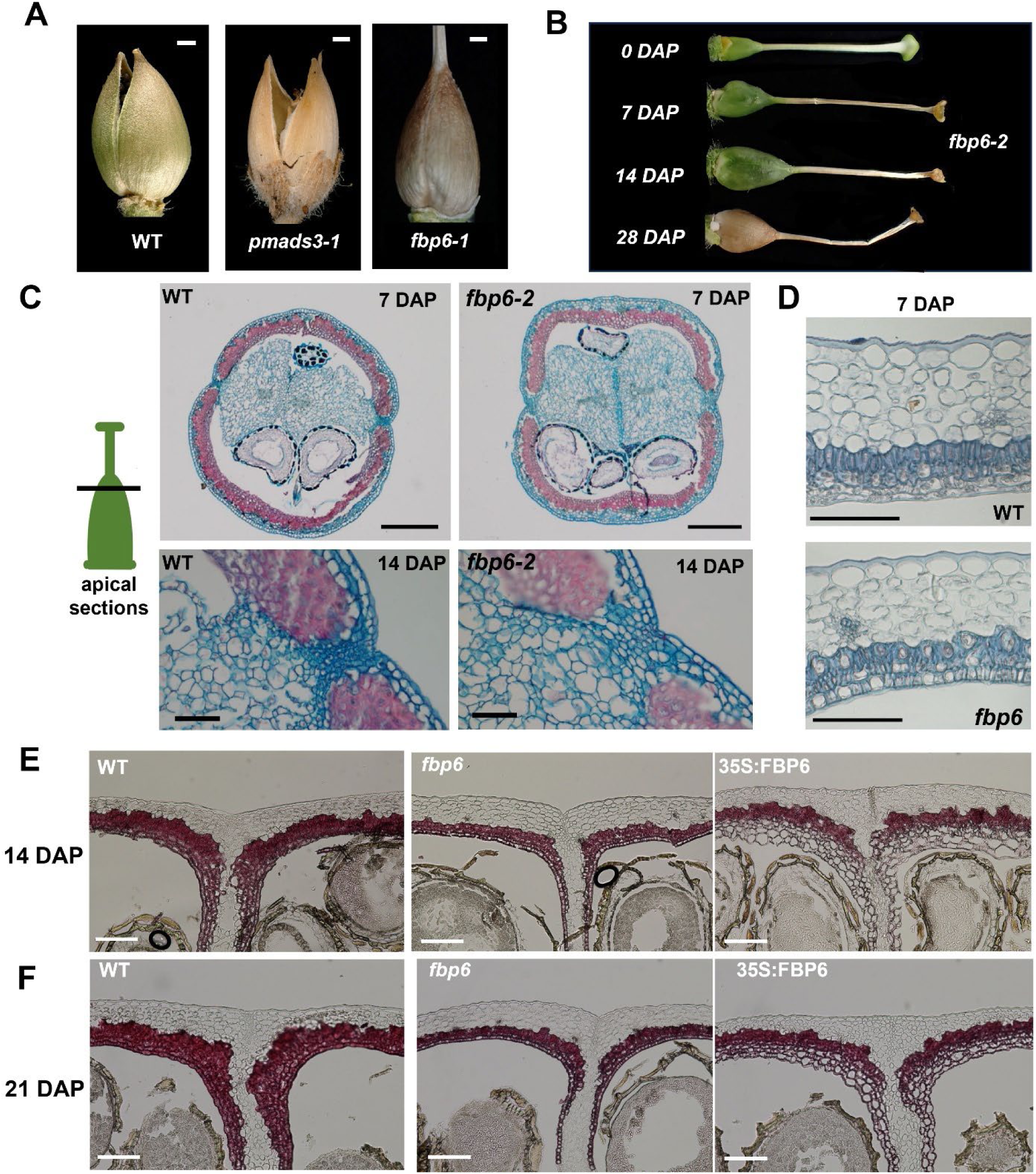
The role of *FBP6* in petunia fruits. [A) macroscopic phenotype of wild-type, *pmads3* and *fbp6* fruits at 5 weeks after pollination. Scale bar = 1mm. B) Series of *fbp6* fruits with the style still attached. C) Phenotype of the *fbp6* mutant fruits in the apical region. Safranin/alcian blue-stained sections from 7 DAP (upper panels) and 14 DAP (lower panels) fruits. The black line in the schematic green ovary depicts the location of the section. D) Toluidine-blue stained pericarp sections of WT and *fbp6* 7 DAP fruits. E-F) Phloroglucinol-stained sections of 14 DAP (E) and 21 DAP (F) fruits at the junction of the two carpels. The phloroglucinol visualizes the lignification in dark red. E) Scale bars upper panels C: 500 µm; scale bars lower panels C: 50 µm; scale bars D-F: 100 µm.

To investigate pericarp development, we sectioned fruits of all lines at 0 DAP, 3 DAP, 7 DAP, 14 DAP and 21 DAP, and stained them with toluidine blue or phloroglucinol. First, we investigated the *fbp6* and *pmads3* single mutant pericarp phenotypes. The *pmads3* mutant fruits largely resemble the wild type, but *fbp6* fruits are different (Fig. 4D-F, Supplementary Figs. S9-11). At the 7 DAP stage, mild differences are observed, with *fbp6* fruits commonly having only three endocarp layers of which the cells appear more regular (Fig. 4D). At later stages, the cells of the innermost endocarp layer are clearly more regular than those of the wild type (opposite to the *quad* phenotype), and all endocarp layers produce significantly less lignin (Fig. 4E-F). In contrast, the *35S:FBP6* fruits at these stages show a phenotype reminiscent of *quad* mutants, with the inner endocarp cells resembling mesocarp cells, being much larger and irregularly shaped (Fig. 4C). While the *quad* endocarp cells produce lignin in a patchy manner, the *35S:FBP6* inner endocarp hardly produces any lignin, not even at 21 DAP (Fig. 4F). Notably, the *35S:pMADS3* fruits show the same phenotype (Supplementary Figs. S11), indicating that the two C-class proteins are functionally interchangeable at the protein level. The *fbp6* phenotype, in combination with the 35S:FBP6 phenotype, suggests that *FBP6* counteracts *PhFUL’s* role in endocarp cell morphology, but may regulate lignin accumulation at a later stage together with the *PhFULs*. While *FBP6* thus appears important for overall pericarp development, the *fbp6* dehiscence failure is most likely attributed to the separation defect in the apical region, particularly because we occasionally observed fruits that were partly cracked open on the sides, but remained attached at the top.

#### The *AP2* co-orthologs *ROB1, ROB2* and *ROB3* redundantly regulate pericarp development

Next, we investigated the role of the *AP2*-like genes in fruit development. From an earlier study (Morel *et al*., 2018), we knew already that ROB1/2/3 together with BEN redundantly and additively act as repressors of nectary development in the apical region of the ovary, with *rob1 rob2 rob3 (rob1/2/3)* and *ben rob1/2/3* mutants typically exhibiting nectaries that strongly extend upwards at the anthesis stage (Morel et al., 2018) (Fig. 5A). This extended nectary covers/replaces the carpel fusion zone where the future fruit normally dehisces, but the effect on fruit dehiscence had not been described (Morel et al., 2018). We found that the extended nectary zone persists throughout fruit development (Fig. 5A) resulting in fruits that completely fail to open to shatter their seeds, and remain firmly closed along the extended nectary. Note that when applying external pressure on *fbp6* dry fruits, these eventually crack open in between the two valves; this is not the case for (*ben) rob1 rob2 rob3* fruits, where seeds can only be released by completely crushing the fruit valves. This indicates that the normal dehiscence zone is entirely absent.

**Figure 5.**
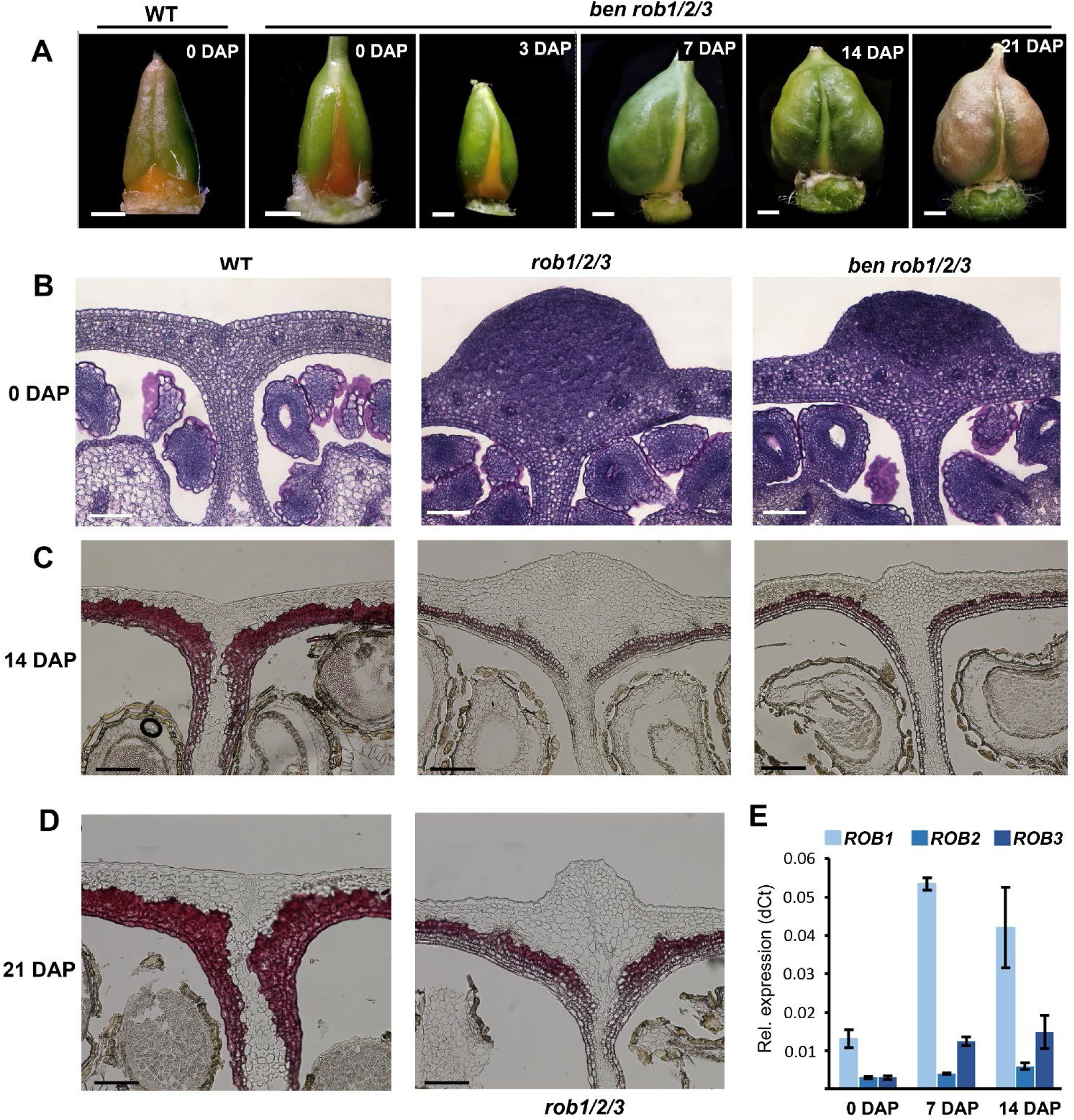
***ROB1*, *ROB2* and *ROB3* redundantly regulate pericarp patterning**. A) macroscopic phenotypes of wild-type at 0 DAP and *ben rob1/2/3* fruits at 0 DAP and later stages. Scale bar = 1mm. B) Toluidine-blue stained pericarp sections of WT and *(ben) rob1/2/3 fruits* at 0 DAP, scale bar = 100 µm; C-D) Phloroglucinol-stained sections of 14 DAP (C) and 21 DAP (D) of WT and *(ben) rob1/2/3* fruits at the junction of the two carpels. Phloroglucinol visualizes the lignification in dark red. Genotypes are indicated above the picture panels (B-C) or below the panels (D). Scale bar (B-D) = 100 µm. E) Relative expression profiles of *ROB1, ROB2* and *ROB3* in 0, 7 and 14 DAP pericarp. Error bars indicate the SE.

Next, we sectioned the fruit at different stages to further investigate the pericarp phenotype. At early stages, there are no clear differences between the WT and *(ben) rob1/2/3* fruits, except for the large nectary bulges (Fig. 5B, and Supplementary Fig. S9). However, at later stages, pericarp patterning seems somewhat delayed, and at 14 DAP, clear differences are apparent (Fig. 5C, Supplementary Fig. S9-S11). At this stage, *rob1/2/3* and *ben rob1/2/3* pericarp contains three layers of small, regular, non-lignified inner endocarp cell layers and only one layer of outer endocarp of which the cells produce some lignin. At 21 DAP, the *rob1/2/3* and *ben rob1/2/3* fruits display much less lignification than the wild type, and have more inner endocarp layers with generally smaller, more regular cells (Fig. 5D). This phenotype suggests that the *AP2*-like genes are required for proper differentiation of all layers, and particularly the outer endocarp layers, which produce most of the lignin. The fact that the cells in the *(ben) rob1/2/3* mutants are more reminiscent of inner-endocarp layer cells may indicate that the AP2-like TFs function opposite to the FUL-like TFs. Because *rob1/2/3* and *ben rob1/2/3* mutants display similar phenotypes, *BEN* does probably not have a major contribution, indicating that fruit patterning is mainly controlled by the three *AP2* co-orthologs. To get more insight into the contributions of the three *ROB* genes to pericarp patterning, we performed expression analysis and found that *ROB1* is considerably higher expressed than *ROB2* and *ROB3* in the gynoecium and at all fruit stages (Fig. 5E, Supplementary Fig. S12). We also determined the fruit phenotypes of several lower order mutants, which revealed that in particular the double mutant *rob1 rob2* displays aberrant endocarp at 7 DAP, similar to *rob1/2/3*. At 14 DAP, the *rob1 rob2* mutant endocarp layers more resemble the wild type, possibly suggesting that *ROB3* is able to compensate for the loss of *ROB1* and *ROB2* on the longer term (Supplementary Fig. S13). In conclusion, the three *ROB* genes redundantly control endocarp development, with *ROB1* probably contributing the most.

### Genetic interactions between the FUL-, SHP-and AP2-like transcription factors

We reasoned that the similarity between the 35S:FBP6 and *quad* phenotypes could be the result of *PhFUL* repression by FBP6. In that case, the 35S:FBP6 overexpression would downregulate the *PhFULs* in the endocarp, leading to large, round endocarp cells. Additionally, the smaller, more regular endocarp cells in the *fbp6* mutant could then be explained by upregulation of the *PhFULs*. To investigate whether there are genetic interactions between the genes in the FUL-SHP-AP2 module, we performed qPCRs to test the expression of *FBP29, FBP26, PFG, euAP1, ROB1, ROB2, ROB3* and *FBP6* in the *quad*, *rob1/2/3*, *fbp6* and WT pericarp (Fig. 6A). All four petunia *AP1/FUL-like* genes were significantly upregulated in the *fbp6* mutant at the 7 DAP and 14 DAP stages, with a particular high upregulation of *FBP29* and *euAP1*. Interestingly, all genes were also upregulated in the *rob1/2/3* mutant, indicating that the petunia AP2 TFs are also repressing the *PhFULs*. Notably, *FBP6* and *pMADS3* are not differentially expressed in the *quad* mutant, but *ROB1* and *ROB3* are upregulated in *quad* fruits at early stages (Figure 6A-C), suggesting that they are repressed by the PhFULs, similar to *AP2* in Arabidopsis (Ripoll *et al*., 2011; Balanzà *et al*., 2018). Additionally, both *ROB1* and *ROB2* are strongly upregulated in the *rob1/2/3* mutant, indicating a negative feedback loop at the *ROB1/2* loci. In conclusion, our data suggest that the *PhFULs* are directly or indirectly repressed by the C-class TFs and AP2-like TFs in the pericarp, while the PhFULs on their turn only repress the *AP2*-like genes, but do not regulate *FBP6*. Together, the interplay between the three TF types appears essential for pericarp patterning, and particularly the establishment and development of the outer and inner endocarp layers.

**Figure 6.**
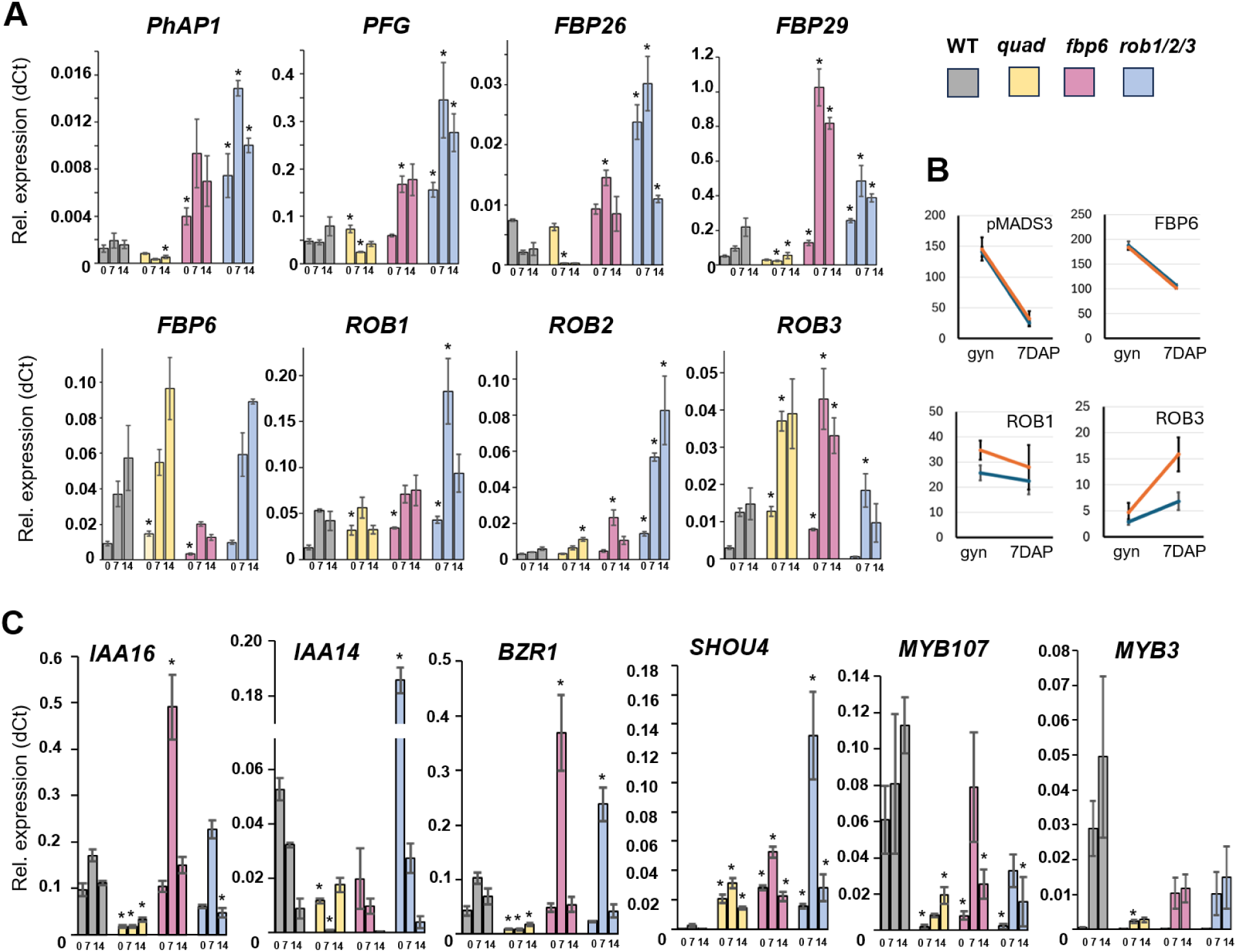
**Genetic interactions between the FUL-, SHP-and AP2-like transcription factors**. A, C) qRT-PCR results of the expression of selected genes in the wild type (grey), *quad* (yellow), *fbp6* (pink) and *rob1/2/3* (blue) mutants at 0 DAP, 7 DAP and 14 DAP in pericarp. B) RPKM plots based on the RNAseq results from a selection of interesting targets from the gynoecium list. Both the expression in the gynoecium and the 7 DAP fruit is depicted. Blue lines indicate the wild-type levels, orange lines the *quad* levels. C) Expression profiles of selected downstream genes. Error bars indicate the SE; significant difference from the wild type at p>0.05 (two-tailed Student T-test) is indicated with an asterisk. (A) expression profiles of the petunia FUL/SHP/AP2-like genes.

### The *FUL*-, *SHP*-, and *AP2*-like genes have overlapping target gene sets

To further study the interactions between the three types of transcription factors, we investigated the expression of selected PhFUL targets in the *fbp6*, *rob1/2/3*, and *quad* mutants by qRT-PCR in 0, 7 and 14 DAP pericarp samples. This unveiled an interesting pattern for the auxin signalling genes, which are downregulated in *quad* pericarp, but upregulated in *fbp6* (*IAA16*-like) and *rob1/2/3* (*IAA14*-like) at 0 DAP. Moreover, the BR-signalling gene *BZR1*-like is upregulated at 7 DAP in both *fbp6* and*rob1/2/3*, while it is downregulated in the *quad* mutant. This suggests that the differentiation of the endocarp is controlled by the opposing effect of PhFUL and ROB/FBP6 on auxin/BR signalling. Interestingly, *SHOU4*-like, the putative negative regulator of cellulose synthases, is strongly upregulated in all three mutants at all stages, with a peak at 7 DAP (Fig. 6C). This may indicate that the entire FUL-AP2-SHP module represses cellulose production to delimit cell layer formation and/or to promote secondary cell wall formation in the endocarp. For the cell-wall modifiers *TCH4*-like, *SKU5*-like, and *IRX15*-like, the differential regulation in *fbp6* or *rob1/2/3* is less pronounced (Supplementary Fig. S14). From 7 DAP onwards, all three mutants show impairment in lignin accumulation. Therefore, we expected that genes related to this phenotype may show similar dysregulation. Indeed, we found that the regulators *MYB107*, *MYB3* and *MYB15* are strongly downregulated in *quad* and also in *fbp6* and *rob1/2/3*, albeit to a lesser extent (Fig. 6C and Supplementary Fig. S14). Thus, the antagonism between PhFUL and ROB/FBP6 acts mainly on the auxin/BZR signalling, while other regulators of (later) pericarp development appear to be regulated in the same direction.

## Discussion

In angiosperms, the ovary starts to develop into a fruit after fertilization. This involves differentiation of the carpel walls (pericarp), which initially consist of layers of undifferentiated parenchyma cells (Pabón-Mora & Litt, 2011). Around 2-4 days after fertilization, different cell layers are specified that support the eventual dispersal of the seeds. In the two main fruit types that are recognized, dry and fleshy fruits, the early differentiation of the pericarp layers occurs similarly, with differentiation into exocarp, mesocarp and endocarp (Pabón-Mora & Litt, 2011; Dardick & Callahan, 2014; Lotz *et al*., 2024). Depending on the dispersal mechanism, additional structures can develop, such as the valve margins in the Arabidopsis silique (Ferrándiz, 2002). At later stages, different metabolic processes are often initiated, such as lignification in dry fruits, or ripening in fleshy fruits. Thus, it appears that early fruit patterning is a conserved process, while development diverges at later stages to give rise to an abundant variation of dry and fleshy fruits. However, it is yet unknown which gene regulatory network could underlie conserved early fruit patterning. Here, we studied in detail the contribution of *FUL*-, *AP2*-and *SHP*-like genes in petunia and propose that these genes may together be part of a conserved module that controls early pericarp differentiation.

### Establishment of the FUL-SHP-AP2 module

Studies in different species have indicated that the transcription factors AP2, SHP and FUL regulate aspects of fruit development in both dry- and fleshy-fruited species (Roeder & Yanofsky, 2006; Vrebalov *et al*., 2009; Jaakola *et al*., 2010; Karlova *et al*., 2011; Ripoll *et al*., 2011; Bemer *et al*., 2012; Pabón-Mora *et al*., 2012; Wang *et al*., 2019; Zhao *et al*., 2019; Chen *et al*., 2020). However, insight in their role in pericarp differentiation is still lacking, as the studies in the model species tomato and Arabidopsis have focused on fruit ripening and the specification of the valve margins, respectively.

We show that in petunia, the FUL-, AP2- and SHP-like transcription factors play an important role at the early stage of fruit development. In the absence of *AP2* activity *(rob1/2/3* mutant), pericarp patterning is delayed, mesocarp cells are smaller, there are more layers with inner-endocarp cells, and very few lignin-producing outer endocarp cells (Fig. 5C-D). This suggests that *AP2* promotes outer endocarp layer identity and possibly also mesocarp identity, resulting in a much stronger inner endocarp establishment in the *rob1/2/3* mutant. In the *quad ap1/ful* mutant, the differentiated endocarp layers are more mesocarp-like (Fig. 1G), suggesting, together with the fact that *PFG* transcripts are localized in the endocarp (Fig. 2E), that the *PhFULs* are required to specify proper endocarp identity. Interestingly, we observed a similar phenotype in fruits of C-class overexpressor lines (35S:FBP6 and 35S:pMADS3), and an opposite phenotype in *fbp6* fruits, pointing to an antagonistic role for *FBP6* in the development of the endocarp layers. Additionally, we show that both the C-class *FBP6* gene and the *AP2-*like *ROB1/2/3* genes repress the petunia *AP1/FULs.* Finally, we found that *FBP6* expression is independent of PhFUL and ROB1/2/3 (Fig. 6A). Altogether, this leads to a model in which the PhFULs promote regular endocarp cells, while FBP6 promotes larger, rounder endocarp cells, which have a mesocarp-like morphology. It is plausible that the PhFUL/FBP6 ratio determines cell identity in the endocarp, with a high PhFUL dosage promoting cell regularity in the inner endocarp, and a higher FBP6 dosage resulting in the more irregular outer-endocarp cells. Also ROB1/2/3 act upstream of the *PhFULs* by repressing *PhFUL* activity and thereby cell regularity. Our data suggest that the PhFULs also repress *ROB1, ROB2* and *ROB3*, which may occur predominantly in the endocarp, where PhFUL activity is high. However, there could also be other factors that lower *ROB1/2/3* transcript accumulation in the endocarp, for example miR172, which degrades *AP2* transcripts in the Arabidopsis inflorescence and fruit (Aukerman & Sakai, 2003; Ripoll *et al*., 2011).

The hypothesis that PhFUL and FBP6/ROB control early endocarp development in an opposite manner is supported by the downstream expression analysis that we performed, which shows that the auxin- and brassinosteroid-signalling genes downstream of the PhFULs are antagonistically regulated by FBP6 and the ROBs. While *IAA14/16* are activated by the PhFULs, *IAA16* is repressed by FBP6 and *IAA14* by ROB1/2/3. Similarly, the brassinosteroid response factor *BZR1* is activated by the PhFULs, but repressed by both FBP6 and the ROBs (Fig. 6C). It appears therefore plausible that the FUL-AP2-SHP module regulates endocarp development via auxin and BR signalling, involving auxin signalling repressors such as *IAA4, IAA14* and *IAA16*, auxin transporters such as *PILS, PIN1* and *PIN3*, and the BR-response factors *BEH1, BEH4* and *BZR1* (Figs. 3 and 6; Supplementary Figure S8). In this early patterning, the transcription factor *NGAL1*, associated with boundary formation (Nicolas *et al*., 2022) and strongly downregulated in the *quad* mutant (Fig. 3C), may play an important role. Interestingly, auxin and BR levels are repressed in boundary zones, associated with reduced differentiation (Wang *et al*., 2016), and PhFULs activation of *IAAs* and *PINs* may facilitate the establishment of a low auxin signalling environment. However, a better spatial and temporal insight in expression levels will be required to support the proposed model, for example by using Laser Capture Microdissection (LCM), single-cell sequencing or Stereo-seq (Xia *et al*., 2022; Li *et al*., 2024).

It remains to be investigated to what extent a FUL-AP2-SHP module acts in other angiosperm species. Given that in Arabidopsis *ful* mutants, the complete conversion to valve margin cells occurs before pericarp layers differentiate (Bowman *et al*., 1999; Roeder & Yanofsky, 2006), it is unclear whether *FUL* is also involved in mesocarp/endocarp differentiation in the Arabidopsis silique. Interestingly, retraction of *FUL* expression away from the replum in *ind alc shp1 shp2* fruits results in mesocarp-like endocarp cells in this region (Liljegren *et al*., 2004; Roeder & Yanofsky, 2006), suggesting that a function in endocarp specification may be conserved. In cucumber, FUL regulates fruit growth partially by repressing auxin transport (Zhao *et al*., 2019), pointing to a more general role of FUL in regulating fruit development via the auxin pathway. Brassinosteroids are common regulators of plant growth and development as well, and regulate for example carpel/silique growth and organ boundary specification in Arabidopsis (Gendron *et al*., 2012; Li *et al*., 2020). Therefore, it would not be surprising if FUL, AP2 and SHP regulate pericarp patterning more generally via the auxin and brassinosteroid pathways, which often act in concert (Tian *et al*., 2018). Future research will have to elucidate this, and additionally reveal whether other fruit factors such as miR172, ARF6/8 (Ripoll *et al*., 2015), RPL (Roeder *et al*., 2003) and IND/HEC3 (Liljegren *et al*., 2004) are part of the module as well. This will particularly require better spatiotemporal studies in early fruit development in different species.

### Additional functions of the petunia *AP1/FUL/SHP* genes in fruit development

In addition to their putative conserved role in early pericarp patterning, our analysis of different order *PhFUL/euAP1* mutants reveals additional functionality both before and after the pericarp patterning stage. Interestingly, we show here that all three petunia *FUL*-like genes contribute to pericarp development, including the euFULII-clade gene *FBP29,* which has not yet been associated with fruit development in other species. Given the early divergence of the euFULII clade, which appears to pre-date the split of the euFULI and AP1 clades (Litt & Irish, 2003; Maheepala *et al*., 2019), this might reflect an ancestral role for the *FUL/AP1* genes in fruit development, although it is also possible that *FBP29* has gained a fruit function in petunia. *EuAP1* has an early role in the restriction of carpel number and placenta growth, and the *PhFULs* contribute to this function, as the multiple-carpel phenotype is only observed in the *quad* mutant and placenta size is larger in the *quad* mutant compared to the *euap1* single mutant. Notably, a multi-carpel phenotype has also been described for fruits of Arabidopsis *pFUL:FUL-VP16* activator lines, which show a stronger mutant phenotype than *ful* mutants (Balanzà *et al*., 2018), as well as for fruits of tomato *ful2* mutants, which regularly have an additional locule (Jiang *et al*., 2021). Besides an early function in carpel/placenta formation, *euAP1* does not appear to contribute to fruit development, in line with its weak expression in the fruit (Fig. 1A), and the role of its orthologs in other species (Pabon-Mora *et al*., 2013).

In the *quad* mutant and triple mutants including *euap1* and *fbp26*, there are more cell-layers at 0 DAP, and this difference remains stable during the early cell division stage. Thus, this phenotype has its origin in the pre-fertilization carpel. Interesting DEGs in the gynoecium that could be involved are the cellulose biosynthesis repressor *SHOU4* and the auxin signalling genes *PILS*, *IAA14* and *IAA16*, because auxin is known to initiate fruit set by activating cell divisions in the pericarp (Dorcey *et al*., 2009; Hu *et al*., 2023). Notably, it has been found that tomato fruits with reduced cytokinin content developed fewer pericarp cell layers, suggesting that the auxin/cytokinin balance influences carpel/pericarp cell layer number (Su *et al*., 2011; Gan *et al*., 2022).

The presence of secondary cell walls is a characteristic of endocarp cells (Dardick & Callahan, 2014), and the fact that we already find important cell wall regulators such as *SHOU4* and *IRX15-L* upregulated in the gynoecium and at 0 DAP (Fig. 3B-C and Fig. 6C), and bound by PFG *in vitro* (Fig. 3E), suggests that the PhFULs may play a direct role in endocarp development and secondary cell wall regulation. While the control of cellulose synthase activity via *SHOU4* (Fig. 6C) (Polko *et al*., 2018) is a shared function of PhFUL, ROB1/2/3 and FBP6, the upregulation of *IRX15-L*, which is responsible for xylan deposition in the secondary cell wall (Brown *et al*., 2011), as well as the downregulation of the putative lignin biosynthesis repressor *MYB3*, is specific for the *quad* mutant (Fig. 6C). Given that lignin deposition in the inner endocarp layers occurs later and is less prominent than in the outer endocarp, this suggests that the PhFULs promote secondary cell wall formation by repressing *SHOU4*, but retard the deposition of secondary metabolites via *IRX15-L* and *MYB3*. This could point to an important role for the PhFULs in inhibiting premature inner endocarp lignification, thereby possibly preventing untimely fruit dehiscence.

FBP6 also contributes to timely fruit dehiscence by promoting the formation of a few cell files with smaller cells at the junction of the two carpels, a function that may be coupled with style abscission (Fig. 4B-C). In addition, the strongly reduced lignification in the *fbp6* mutant points to a role for *FBP6* in the promotion of lignification in the outer endocarp layers. These results are in agreement with the phenotype of NbSHP-VIGS plants in Nicotiana, which exhibit non-dehiscent fruits with reduced lignification (Fourquin & Ferrándiz, 2012).

Possibly, the combined action of *FBP6* and the *PhFULs* at later stages results in coordinated lignification in the inner and outer endocarp to induce carpel rupture at the location of the small cell files, followed by fruit opening and seed shattering. Such a joint function may be conserved in Arabidopsis, where the lignification in the endocarp *b* layer is only lost in quintuple *ful shp1 shp2 ind alc* mutants (Liljegren *et al*., 2004; Roeder & Yanofsky, 2006).

The phenotype of the *rob1/2/3* mutant fruits appears more severe than that of the *fbp6* and *quad* fruits, and it is difficult to uncouple different phenotypic aspects. The strongly extended nectaries that cover/replace the carpel fusion zone may prevent dehiscence, but failure of dehiscence may also be due to the very limited lignification in the endocarp layers. It is clear however, that the *AP2* orthologs are essential for a proper establishment of the endocarp layers.

In conclusion, using the model petunia to investigate the role of the FUL-AP2-SHP module in fruit development, we gained insight in a possible conserved pericarp-patterning network. It appears plausible that there is a conserved network acting during early fruit development, while it diverges at later stages. We hypothesize that, similar to the conserved ABC-model that defines the identity of the different floral organs, there may be a conserved module that defines early pericarp development, and includes FUL, AP2 and SHP. However, to validate this model in petunia and other species, more studies, with in particular detailed spatiotemporal expression analysis, are required. New technologies based on spatial transcriptomics are particularly suited for these kind of studies, but are still quite expensive.

## Supplementary data

**Figure S1.** Safranin/alcian blue stained sections of fruits of WT and different double mutant combinations at 14 DAP.

**Figure S2**. Safranin/alcian blue stained sections of fruits of WT, *FUL/AP1* quadruple mutant, and different triple mutant combinations showing the differences in placenta size.

**Figure S3.** Safranin/alcian blue stained sections of fruits of WT and *FUL/AP1* quadruple mutant fruits at different time points after pollination.

**Figure S4.** Stained sections of fruits of WT and *FUL/AP1* triple mutant fruits at different time points after pollination.

**Figure S5**. Safranin/alcian blue-stained sections from 14 DAP fruits of different mutant combinations of the petunia *FUL*-like triples and the WT.

**Figure S6.** In situ hybridisations of WT fruits at 3, 5 and 7 DAP.

**Figure S7**. Results of a principal component analysis (PCA).

**Figure S8**. RPKM plots of DEGs in the *quad-ful* mutant pericarp in the gynoecium and at 7 DAP.

**Figure S9.** Toluidine blue-stained sections of the ovary wall of early fruit stages of WT and mutant combinations of petunia *FUL*-like, *SHP*-like and *AP2*-like.

**Figure S10.** Toluidine blue-stained sections from 7 DAP fruits of different mutant combinations of petunia *FUL*-like, *SHP*-like and *AP2*-like and the WT.

**Figure S11.** Phloroglucinol-stained sections from 14 DAP fruits of different mutant combinations of petunia *FUL*-like, *SHP*-like and *AP2*-like and the WT.

**Figure S12.** RPKM plots of the petunia *FUL-, AP2-*and *SHP*-like genes at 0 DAP and 7 DAP.

**Figure S13.** Safranin/alcian-blue-stained sections of different mutant combinations of *ben, rob1, rob2* and *rob3* at 7 DAP.

**Figure S14.** Expression data of selected targets in WT, *quad, fbp6* and *rob1/2/3*.

**Table S1.** Oligo sequences used in this study **Table S2.** Read count data and DEGs at 7 DAP. **Table S3.** Read count data and DEGS gynoecium.

**Table S4.** Overlapping DEGs in the gynoecium and 7 DAP samples.

**Table S5**. GO term enrichment analysis for the gynoecium and 7 DAP samples.

**Table S6.** Interesting DEGs in gynoecium and at 7 DAP.

## Acknowledgements

M.V wishes to thank D. Lemaitre for her help on the project during her internship in the team, and the RDP plant culture team for their help in taking care of the petunia plants. We also appreciate the help of Robin van Velzen in uploading the high-throughput sequencing data to ENA.

## Author contributions

MB and MV conceptualization; MB, ER, CG-M, NHC, PM, PC and MV: methodology; PM, MB and MV: formal analysis; MB, PC, NHC, PM, ER, KT, EA, SRB and MV: investigation; MB, CG-M and MV: resources; MB: data curation; MB and MV: writing – original draft; MB, KT, PM and MV: writing – review & editing; MB, PC, NHC, PM, ER and MV: visualization; MB and MV: supervision. MB, CG-M and MV: funding acquisition.

## Data availability

The raw data of the RNAseq experiment has been deposited at the European Nucleotide Archive (ENA, https://ena-docs.readthedocs.io/en/latest/retrieval/general-guide.html) under project PRJEB99017.

## References

Aerts N, de Bruijn S, van Mourik H, Angenent GC, van Dijk ADJ. 2018. Comparative analysis of binding patterns of MADS-domain proteins in Arabidopsis thaliana. BMC Plant Biol 18(1): 131.

Aukerman MJ, Sakai H. 2003. Regulation of flowering time and floral organ identity by a MicroRNA and its APETALA2-like target genes. Plant Cell 15(11): 2730-2741.

Balanzà V, Martínez-Fernández I, Sato S, Yanofsky MF, Kaufmann K, Angenent GC, Bemer M, Ferrándiz C. 2018. Genetic control of meristem arrest and life span in Arabidopsis by a FRUITFULL-APETALA2 pathway. Nat Commun 9(1): 565.

Barbez E, Kubeš M, Rolčík J, Béziat C, Pěnčík A, Wang B, Rosquete MR, Zhu J, Dobrev PI, Lee Y, et al. 2012. A novel putative auxin carrier family regulates intracellular auxin homeostasis in plants. Nature 485(7396): 119-122.

Bemer M, Karlova R, Ballester AR, Tikunov YM, Bovy AG, Wolters-Arts M, Rossetto Pde B, Angenent GC, de Maagd RA. 2012. The tomato FRUITFULL homologs TDR4/FUL1 and MBP7/FUL2 regulate ethylene-independent aspects of fruit ripening. Plant Cell 24(11): 4437-4451.

Bemer M, van Mourik H, Muiño JM, Ferrándiz C, Kaufmann K, Angenent GC. 2017. FRUITFULL controls SAUR10 expression and regulates Arabidopsis growth and architecture. J Exp Bot 68(13): 3391-3403.

Bombarely A, Moser M, Amrad A, Bao M, Bapaume L, Barry CS, Bliek M, Boersma MR, Borghi L, Bruggmann R, et al. 2016. Insight into the evolution of the Solanaceae from the parental genomes of Petunia hybrida. Nature Plants 2(6): 16074.

Bowman JL, Baum SF, Eshed Y, Putterill J, Alvarez J. 1999. Molecular genetics of gynoecium development in Arabidopsis. Curr Top Dev Biol 45: 155-205.

Bowman JL, Moyroud E. 2024. Reflections on the ABC model of flower development. The Plant Cell 36(5): 1334-1357.

Braatz J, Harloff HJ, Mascher M, Stein N, Himmelbach A, Jung C. 2017. CRISPR-Cas9 Targeted Mutagenesis Leads to Simultaneous Modification of Different Homoeologous Gene Copies in Polyploid Oilseed Rape (Brassica napus). Plant Physiol 174(2): 935-942.

Brown D, Wightman R, Zhang Z, Gomez LD, Atanassov I, Bukowski J-P, Tryfona T, McQueen-Mason SJ, Dupree P, Turner S. 2011. Arabidopsis genes IRREGULAR XYLEM (IRX15) and IRX15L encode DUF579-containing proteins that are essential for normal xylan deposition in the secondary cell wall. The Plant Journal 66(3): 401-413.

Cartolano M, Castillo R, Efremova N, Kuckenberg M, Zethof J, Gerats T, Schwarz-Sommer Z, Vandenbussche M. 2007. A conserved microRNA module exerts homeotic control over Petunia hybrida and Antirrhinum majus floral organ identity. Nat Genet 39(7): 901-905.

Chen C, Zhang Y, Cai J, Qiu Y, Li L, Gao C, Gao Y, Ke M, Wu S, Wei C, et al. 2023. Multi-copper oxidases SKU5 and SKS1 coordinate cell wall formation using apoplastic redox-based reactions in roots. Plant Physiol 192(3): 2243-2260.

Chen L-G, Gao Z, Zhao Z, Liu X, Li Y, Zhang Y, Liu X, Sun Y, Tang W. 2019. BZR1 Family Transcription Factors Function Redundantly and Indispensably in BR Signaling but Exhibit BRI1-Independent Function in Regulating Anther Development in Arabidopsis. Molecular Plant 12(10): 1408-1415.

Chen T, Qin G, Tian S. 2020. Regulatory network of fruit ripening: current understanding and future challenges. New Phytologist 228(4): 1219-1226.

Chezem WR, Memon A, Li FS, Weng JK, Clay NK. 2017. SG2-Type R2R3-MYB Transcription Factor MYB15 Controls Defense-Induced Lignification and Basal Immunity in Arabidopsis. Plant Cell 29(8): 1907-1926.

Choi SJ, Lee Z, Kim S, Jeong E, Shim JS. 2023. Modulation of lignin biosynthesis for drought tolerance in plants. Front Plant Sci 14: 1116426.

Chung M-Y, Vrebalov J, Alba R, Lee J, McQuinn R, Chung J-D, Klein P, Giovannoni J. 2010. A tomato (Solanum lycopersicum) APETALA2/ERF gene, SlAP2a, is a negative regulator of fruit ripening. The Plant Journal 64(6): 936–947.

Dardick C, Callahan AM. 2014. Evolution of the fruit endocarp: molecular mechanisms underlying adaptations in seed protection and dispersal strategies. Front Plant Sci 5: 284.

Dorcey E, Urbez C, Blázquez MA, Carbonell J, Perez-Amador MA. 2009. Fertilization-dependent auxin response in ovules triggers fruit development through the modulation of gibberellin metabolism in Arabidopsis. Plant J 58(2): 318-332.

Ferrándiz C. 2002. Regulation of fruit dehiscence in Arabidopsis. J Exp Bot 53(377): 2031-2038.

Ferrándiz C, Liljegren SJ, Yanofsky MF. 2000. Negative regulation of the SHATTERPROOF genes by FRUITFULL during Arabidopsis fruit development. Science 289(5478): 436-438.

Ferrario S. 2004. Functional characterization of MADS box transcription factors in Petunia hybrida. *PhD thesis Radboud University* https://edepot.wur.nl/23355.

Flütsch S, Nigro A, Conci F, Fajkus J, Thalmann M, Trtílek M, Panzarová K, Santelia D. 2020. Glucose uptake to guard cells via STP transporters provides carbon sources for stomatal opening and plant growth. EMBO Rep 21(8): e49719.

Fourquin C, Ferrándiz C. 2012. Functional analyses of AGAMOUS family members in Nicotiana benthamiana clarify the evolution of early and late roles of C-function genes in eudicots. The Plant Journal 71(6): 990-1001.

Gan L, Song M, Wang X, Yang N, Li H, Liu X, Li Y. 2022. Cytokinins are involved in regulation of tomato pericarp thickness and fruit size. Horticulture research 9: uhab041.

Gendron JM, Liu J-S, Fan M, Bai M-Y, Wenkel S, Springer PS, Barton MK, Wang Z-Y. 2012. Brassinosteroids regulate organ boundary formation in the shoot apical meristem of Arabidopsis. Proceedings of the National Academy of Sciences 109(51): 21152-21157.

Geng P, Zhang S, Liu J, Zhao C, Wu J, Cao Y, Fu C, Han X, He H, Zhao Q. 2020. MYB20, MYB42, MYB43, and MYB85 Regulate Phenylalanine and Lignin Biosynthesis during Secondary Cell Wall Formation. Plant Physiol 182(3): 1272–1283.

Gimenez E, Castañeda L, Pineda B, Pan IL, Moreno V, Angosto T, Lozano R. 2016. TOMATO AGAMOUS1 and ARLEQUIN/TOMATO AGAMOUS-LIKE1 MADS-box genes have redundant and divergent functions required for tomato reproductive development. Plant Molecular Biology 91(4): 513-531.

Gómez-Mena C, Roque EM. 2018. Non-isotopic RNA In Situ Hybridization for Functional Analyses in Medicago truncatula. Methods Mol Biol 1822: 133-144.

Gu Q, Ferrandiz C, Yanofsky MF, Martienssen R. 1998a. The FRUITFULL MADS-box gene mediates cell differentiation during Arabidopsis fruit development. Development 125(8): 1509.

Gu Q, Ferrándiz C, Yanofsky MF, Martienssen R. 1998b. The FRUITFULL MADS-box gene mediates cell differentiation during Arabidopsis fruit development. Development 125(8): 1509-1517.

Heijmans K, Ament K, Rijpkema AS, Zethof J, Wolters-Arts M, Gerats T, Vandenbussche M. 2012a. Redefining C and D in the Petunia ABC. The Plant Cell 24(6): 2305.

Heijmans K, Morel P, Vandenbussche M. 2012b. MADS-box Genes and Floral Development: the Dark Side. Journal of Experimental Botany 63(15): 5397-5404.

Hu J, Li X, Sun TP. 2023. Four class A AUXIN RESPONSE FACTORs promote tomato fruit growth despite suppressing fruit set. Nat Plants 9(5): 706-719.

Immink RG, Ferrario S, Busscher-Lange J, Kooiker M, Busscher M, Angenent GC. 2003. Analysis of the petunia MADS-box transcription factor family. Mol Genet Genomics 268(5): 598-606.

Jaakola L, Poole M, Jones MO, Kämäräinen-Karppinen T, Koskimäki JJ, Hohtola A, Häggman H, Fraser PD, Manning K, King GJ, et al. 2010. A SQUAMOSA MADS Box Gene Involved in the Regulation of Anthocyanin Accumulation in Bilberry Fruits Plant Physiol153(4): 1619–1629.

Jiang X, Lubini G, Hernandes-Lopes J, Rijnsburger K, Veltkamp V, de Maagd RA, Angenent GC, Bemer M. 2021. FRUITFULL-like genes regulate flowering time and inflorescence architecture in tomato. The Plant Cell 34(3): 1002-1019.

Johansen DE. 1941. Plant Microtechnique. Nature 147(3721): 222-222.

Karlova R, Rosin FM, Busscher-Lange J, Parapunova V, Do PT, Fernie AR, Fraser PD, Baxter C, Angenent GC, de Maagd RA. 2011. Transcriptome and Metabolite Profiling Show That APETALA2a Is a Major Regulator of Tomato Fruit Ripening. The Plant Cell 23(3): 923.

Li T, Kang X, Wei L, Zhang D, Lin H. 2020. A gain-of-function mutation in Brassinosteroid-insensitive 2 alters Arabidopsis floral organ development by altering auxin levels. Plant Cell Rep 39(2): 259-271.

Li X, Li B, Gu S, Pang X, Mason P, Yuan J, Jia J, Sun J, Zhao C, Henry R. 2024. Single-cell and spatial RNA sequencing reveal the spatiotemporal trajectories of fruit senescence. Nat Commun 15(1): 3108.

Liljegren SJ, Ditta GS, Eshed Y, Savidge B, Bowman JL, Yanofsky MF. 2000. SHATTERPROOF MADS-box genes control seed dispersal in Arabidopsis. Nature 404(6779): 766-770.

Liljegren SJ, Roeder AH, Kempin SA, Gremski K, Østergaard L, Guimil S, Reyes DK, Yanofsky MF. 2004. Control of fruit patterning in Arabidopsis by INDEHISCENT. Cell 116(6): 843-853.

Litt A, Irish VF. 2003. Duplication and diversification in the APETALA1/FRUITFULL floral homeotic gene lineage: implications for the evolution of floral development. Genetics 165(2): 821-833.

Lotz D, Rössner LH, Ehlers K, Kong D, Rössner C, Rupp O, Becker A. 2024. Conservation of the dehiscence zone gene regulatory network in dicots and the role of the SEEDSTICK ortholog of California poppy (Eschscholzia californica) in fruit development. Evodevo 15(1): 16.

Maheepala DC, Emerling CA, Rajewski A, Macon J, Strahl M, Pabón-Mora N, Litt A. 2019. Evolution and Diversification of FRUITFULL Genes in Solanaceae. Front Plant Sci 10: 43.

Metsalu T, Vilo J. 2015. ClustVis: a web tool for visualizing clustering of multivariate data using Principal Component Analysis and heatmap. Nucleic Acids Res 43(W1): W566-W570.

Mishra BS, Sharma M, Laxmi A. 2022. Role of sugar and auxin crosstalk in plant growth and development. Physiologia Plantarum 174(1): e13546.

Morel P, Chambrier P, Boltz V, Chamot S, Rozier F, Rodrigues Bento S, Trehin C, Monniaux M, Zethof J, Vandenbussche M. 2019. Divergent Functional Diversification Patterns in the SEP/AGL6/AP1 MADS-Box Transcription Factor Superclade. Plant Cell 31(12): 3033-3056.

Morel P, Heijmans K, Ament K, Chopy M, Trehin C, Chambrier P, Rodrigues Bento S, Bimbo A, Vandenbussche M. 2018. The Floral C-Lineage Genes Trigger Nectary Development in Petunia and Arabidopsis. Plant Cell 30(9): 2020-2037.

Morel P, Heijmans K, Rozier F, Zethof J, Chamot S, Bento SR, Vialette-Guiraud A, Chambrier P, Trehin C, Vandenbussche M. 2017. Divergence of the Floral A-Function between an Asterid and a Rosid Species. Plant Cell 29(7): 1605-1621.

Moyroud E, Glover BJ. 2017. The Evolution of Diverse Floral Morphologies. Current Biology 27(17): R941-R951.

Nicolas A, Maugarny-Calès A, Adroher B, Chelysheva L, Li Y, Burguet J, Bågman AM, Smit ME, Brady SM, Li Y, et al. 2022. De novo stem cell establishment in meristems requires repression of organ boundary cell fate. Plant Cell 34(12): 4738-4759.

Nookaraju A, Pandey SK, Ahlawat YK, Joshi CP. 2022. Understanding the Modus Operandi of Class II KNOX Transcription Factors in Secondary Cell Wall Biosynthesis. Plants (Basel) 11(4).

Østergaard L, Kempin SA, Bies D, Klee HJ, Yanofsky MF. 2006. Pod shatter-resistant Brassica fruit produced by ectopic expression of the FRUITFULL gene. Plant Biotechnol J 4(1): 45-51.

Pabón-Mora N, Ambrose BA, Litt A. 2012. Poppy APETALA1/FRUITFULL orthologs control flowering time, branching, perianth identity, and fruit development. Plant Physiol 158(4): 1685-1704.

Pabon-Mora N, Hidalgo O, Gleissberg S, Litt A. 2013. Assessing duplication and loss of APETALA1/FRUITFULL homologs in Ranunculales. Front Plant Sci Volume 4 -2013.

Pabón-Mora N, Litt A. 2011. Comparative anatomical and developmental analysis of dry and fleshy fruits of Solanaceae. Am J Bot 98(9): 1415-1436.

Polko JK, Barnes WJ, Voiniciuc C, Doctor S, Steinwand B, Hill JL, Jr., Tien M, Pauly M, Anderson CT, Kieber JJ. 2018. SHOU4 Proteins Regulate Trafficking of Cellulose Synthase Complexes to the Plasma Membrane. Current Biology 28(19): 3174-3182.e3176.

Qin W, Yin Q, Chen J, Zhao X, Yue F, He J, Yang L, Liu L, Zeng Q, Lu F, et al. 2020. The class II KNOX transcription factors KNAT3 and KNAT7 synergistically regulate monolignol biosynthesis in Arabidopsis. J Exp Bot 71(18): 5469-5483.

Quinn O, Kumar M, Turner S. 2024. The role of lipid-modified proteins in cell wall synthesis and signaling. Plant Physiol 194(1): 51-66.

Rajani S, Sundaresan V. 2001. The Arabidopsis myc/bHLH gene ALCATRAZ enables cell separation in fruit dehiscence. Curr Biol 11(24): 1914-1922.

Ripoll JJ, Bailey LJ, Mai Q-A, Wu SL, Hon CT, Chapman EJ, Ditta GS, Estelle M, Yanofsky MF. 2015. microRNA regulation of fruit growth. Nature Plants 1(4): 15036.

Ripoll JJ, Roeder AH, Ditta GS, Yanofsky MF. 2011. A novel role for the floral homeotic gene APETALA2 during Arabidopsis fruit development. Development 138(23): 5167-5176.

Roeder AH, Ferrándiz C, Yanofsky MF. 2003. The role of the REPLUMLESS homeodomain protein in patterning the Arabidopsis fruit. Curr Biol 13(18): 1630-1635.

Roeder AH, Yanofsky MF. 2006. Fruit development in Arabidopsis. Arabidopsis Book 4: e0075.

Shan H, Cheng J, Zhang R, Yao X, Kong H. 2019. Developmental mechanisms involved in the diversification of flowers. Nature Plants 5(9): 917-923.

Shi H, Li X, Lv M, Li J. 2022. BES1/BZR1 Family Transcription Factors Regulate Plant Development via Brassinosteroid-Dependent and Independent Pathways. Int J Mol Sci 23(17).

Smaczniak C, Immink RG, Muiño JM, Blanvillain R, Busscher M, Busscher-Lange J, Dinh QD, Liu S, Westphal AH, Boeren S, et al. 2012. Characterization of MADS-domain transcription factor complexes in Arabidopsis flower development. Proc Natl Acad Sci U S A 109(5): 1560-1565.

Smaczniak C, Immink RGH, Angenent GC, Kaufmann K. 2012. Developmental and evolutionary diversity of plant MADS-domain factors: insights from recent studies. Development 139(17): 3081.

Su Y-H, Liu Y-B, Zhang X-S. 2011. Auxin–Cytokinin Interaction Regulates Meristem Development. Molecular Plant 4(4): 616-625.

Sun L, Feraru E, Feraru MI, Waidmann S, Wang W, Passaia G, Wang Z-Y, Wabnik K, Kleine-Vehn J. 2020. PIN-LIKES Coordinate Brassinosteroid Signaling with Nuclear Auxin Input in Arabidopsis thaliana. Current Biology 30(9): 1579–1588.e1576.

Thoris K, Correa Marrero M, Fiers M, Lai X, Zahn IE, Jiang X, Mekken M, Busscher S, Jansma S, Nanao M, et al. 2024. Uncoupling FRUITFULL’s functions through modification of a protein motif identified by co-ortholog analysis. Nucleic Acids Res 52(21): 13290-13304.

Tian H, Lv B, Ding T, Bai M, Ding Z. 2018. Auxin-BR Interaction Regulates Plant Growth and Development. Front Plant Sci Volume 8 -2017.

van Mourik H, Chen P, Smaczniak C, Boeren S, Kaufmann K, Bemer M, Angenent GC, Muino JM. 2023. Dual specificity and target gene selection by the MADS-domain protein FRUITFULL. Nat Plants 9(3): 473-485.

Vandenbussche M, Janssen A, Zethof J, Van Orsouw N, Peters J, Van Eijk MJT, Rijpkema AS, Schneiders H, Santhanam P, De Been M, et al. 2008. Generation of a 3D indexed Petunia insertion database for reverse genetics. The Plant Journal 54(6): 1105-1114.

Vandenbussche M, Zethof J, Souer E, Koes R, Tornielli GB, Pezzotti M, Ferrario S, Angenent GC, Gerats T. 2003. Toward the analysis of the petunia MADS box gene family by reverse and forward transposon insertion mutagenesis approaches: B, C, and D floral organ identity functions require SEPALLATA-like MADS box genes in petunia. Plant Cell 15(11): 2680-2693.

Vrebalov J, Pan IL, Arroyo AJM, McQuinn R, Chung M, Poole M, Rose J, Seymour G, Grandillo S, Giovannoni J, et al. 2009. Fleshy Fruit Expansion and Ripening Are Regulated by the Tomato <em>SHATTERPROOF</em> Gene <em>TAGL1</em&gt. The Plant Cell 21(10): 3041.

Wang Q, Hasson A, Rossmann S, Theres K. 2016. Divide et impera: boundaries shape the plant body and initiate new meristems. New Phytologist 209(2): 485-498.

Wang R, Tavano ECdR, Lammers M, Martinelli AP, Angenent GC, de Maagd RA. 2019. Re-evaluation of transcription factor function in tomato fruit development and ripening with CRISPR/Cas9-mutagenesis. Scientific Reports 9(1): 1696.

Wang S, Yamaguchi M, Grienenberger E, Martone PT, Samuels AL, Mansfield SD. 2020. The Class II KNOX genes KNAT3 and KNAT7 work cooperatively to influence deposition of secondary cell walls that provide mechanical support to Arabidopsis stems. Plant J 101(2): 293-309.

Woolfson KN, Esfandiari M, Bernards MA. 2022. Suberin Biosynthesis, Assembly, and Regulation. Plants (Basel) 11(4).

Xia K, Sun HX, Li J, Li J, Zhao Y, Chen L, Qin C, Chen R, Chen Z, Liu G, et al. 2022. The single-cell stereo-seq reveals region-specific cell subtypes and transcriptome profiling in Arabidopsis leaves. Dev Cell 57(10): 1299-1310.e1294.

Zhai Y, Cai S, Hu L, Yang Y, Amoo O, Fan C, Zhou Y. 2019. CRISPR/Cas9-mediated genome editing reveals differences in the contribution of INDEHISCENT homologues to pod shatter resistance in Brassica napus L. Theor Appl Genet 132(7): 2111-2123.

Zhang C, He M, Jiang Z, Liu L, Pu J, Zhang W, Wang S, Xu F. 2022. The Xyloglucan Endotransglucosylase/Hydrolase Gene XTH22/TCH4 Regulates Plant Growth by Disrupting the Cell Wall Homeostasis in Arabidopsis under Boron Deficiency. International Journal of Molecular Sciences 10.3390/ijms23031250

Zhao J, Jiang L, Che G, Pan Y, Li Y, Hou Y, Zhao W, Zhong Y, Ding L, Yan S, et al. 2019. A Functional Allele of CsFUL1 Regulates Fruit Length through Repressing CsSUP and Inhibiting Auxin Transport in Cucumber. Plant Cell 31(6): 1289-1307.

Zhou J, Lee C, Zhong R, Ye ZH. 2009. MYB58 and MYB63 are transcriptional activators of the lignin biosynthetic pathway during secondary cell wall formation in Arabidopsis. Plant Cell 21(1): 248-266.

Zhou K. 2019. GPI-anchored SKS proteins regulate root development through controlling cell polar expansion and cell wall synthesis. Biochemical and Biophysical Research Communications 509(1): 119-124.

Zhou M, Zhang K, Sun Z, Yan M, Chen C, Zhang X, Tang Y, Wu Y. 2017. LNK1 and LNK2 Corepressors Interact with the MYB3 Transcription Factor in Phenylpropanoid Biosynthesis. Plant Physiol 174(3): 1348-1358.

Zumajo-Cardona C, Ambrose BA, Madrigal Y, Pabón-Mora N. 2025. Dehiscent fruits in Brassicaceae and Papaveraceae: convergent morpho-anatomical features with divergent underlying genetic mechanisms. Ann Bot 136(7): 1429-1439.

