## Supplemental Table 1 for "A FUL-SHP-AP2 module regulates fruit development in petunia"

**Supplemental Table 6:** Oligo Sequences Used in this Study.

| **Name** | **Forward primer (5’-3’)** | **Reverse primer (5’-3’)** |
| --- | --- | --- |
| **Genotyping** | | |
| **Ph-euAP1-317** | CAGGAGAACTGGAGTCAGCTG | GAAACAAGCATAACATTCCTGAATGG |
| **PFG-11** | TTTATTCTGCACTATACTTTTTTGTTAGAC | CGACGTTTTGAAAAAGTAACTTGTC |
| **FBP26-76** | CATACCCTAGTGCAGGAGAC | CAGCATCACAAAGCACAGAAATC |
| **FBP29-153** | CCTTCTCTAAGAGACGCTCTGGATTG | GAGTACTCAAATAACTTGCCTTTGGTA |
| **FBP6-155** | ATGGTGTTTCCTAATCAAGAATTTGAG | CAACTTCAGCATCACAGAGCAC |
| **PMADS3-1** | ATTGCAAGAGAGATCTCTCCACAA | AAACTTGCGTCTCTTGCAAAAAGT |
| **BEN-724** | AGAGTTTGTGCATATACTTCGACG | CAGGCAACTCACTTCTTTCCAAG |
| **ROB1-61** | GGGATCTAAATGATTCTCCAGATC | GCTATTGCTGATGAACTTGAAGTC |
| **ROB2-915** | TTATTAGTGAGGTTGAGCCTTGAG | CTACGATCAAAGTTAGTAACTGC |
| **ROB3-935** | CTTAGCGTTCTTTTTAACTACTGAAC | CTGGAATCGAAGTTAGTGACTG |
| **In situ hybridisation** | | |
|  | F (sense) | R (antisense) |
| PFG | TAATACGACTCACTATAGGAAATCTTCTAGGGCCAGCA | TAATACGACTCACTATAGGAAATCTTCTAGGGCCAGCA |
| FBP29 | TATTTAGGTGACACTATAGAAGTGGGCAAACACAACC | TAATACGACTCACTATAGCAATGGGTTCCAACTTCC |
| FBP26 | TATTTAGGTGACACTATAGCAGAGTCAGTGGGAACCA | TAATACGACTCACTATAGCACATGATATTGTATATGTAC |
| **qRT- PCR analysis** | | |
| ***RAN*** | AAGCTCCCACCTGTCTGGAAA | AACAGATTGCCGGAAGCCA |
| ***GAPDH*** | ACTTTGTTGGTGACAGCAGGT | TCATACCATGACACAACTTTCACA |
| ***FBP26*** | CACAGCCCTTGAACTCTCTT | GAGATGGCGAAGCATCCAT |
| ***FBP29*** | TACCTCAGCCTCCATCACTATC | TGGCGGCATCTGAGAATTT |
| ***PFG*** | CCATTGGACTCTCCTCACCTA | GCTGTGAAGCTCCTTCTACTTC |
| ***Ph-euAP1*** | ATGCGTCAATCTCCGAACTAC | TTGCTGCCTTACTGTCTTATCC |
| ***ROB1*** | TGAGACCTGGTGAATCCCATA | CTGCCGCTTGGAAATTGA |
| ***ROB2*** | CCTCGACAGTTCAGCTCGTA | GTGAAAGCACATCTCTTGCATT |
| ***ROB3*** | TGGGAAGGATGCAGTCACTA | GCTGTGATCTAATGCCTTATGAGTAG |
| ***FBP6*** | CGTCTCTATGAATATGCCAACAACAG | TGCTCAATGCCTCTCCAACA |
| ***FBP13*** | AGTTGAGGATGATTGCTCAGATA | TTGCATGAAGTGTCATCACAAA |
| ***FBP21*** | GGAACAGATTGCACGACTAAAAG | CCTGCTCTCCACTTGATACTTG |
| ***Peaxi162Scf00111g00022*** | CCGCAGCATTTCACACTTAC | CCATTGCTTTCGTGGTTATTGA |
| ***Peaxi162Scf00385g00415*** | TTGGTGCAACATGATCCATTATATC | CTCTCACCAGCTCCAAATAACT |
| ***Peaxi162Scf00442g00524*** | CTCGAGTGATGACCGCATAATG | CATTTTCATGGCGCCTTACC |
| ***Peaxi162Scf01181g00022*** | GTGGAGTCCTGAGAAGAGAAAG | CTGCCCATGAGTTAGGATAGAC |
| ***Peaxi162Scf00549g00028*** | ATGAATGCAGGCCAGTACAA | CCCTCTTGGACCATCAACTAAA |
| ***Peaxi162Scf00006g00098*** | ATTTGAGGAATCAAGTGGAATAAGC | GAGATACAAGCCCATGGTCAA |
| ***Peaxi162Scf00007g02122*** | TCCTCGTTTAACTTAGATGATTACCTC | GTGCTCATCTCTAACACCCAAG |
| ***Peaxi162Scf00112g01217*** | GCACCACTTATAGAAAGGGATG | GCTCATTTGGTAAGACGAACAAG |
| ***Peaxi162Scf00305g00045*** | AGGACTCTTATCAGAATATGAAGAG | GGCAATCTTTGACCACCTGT |
| ***Peaxi162Scf00543g00414*** | GGCTGAGGACTTATTAGACTTACC | ACCCAGCACCAGTTTAGTTAG |
| ***Peaxi162Scf00572g00812*** | TGAGAGGAACTGAAGCAATAGG | CTCGGTTCACCTTCTCGAATA |
| ***Peaxi162Scf01327g00017*** | TGATCTAGAGCTCACTCTTGGA | GCATAAGAGGTCCTACATATTCCTATC |
| ***Peaxi162Scf01416g00118*** | AGGATCAGAAGCTATTGGACTTG | CATGCTTTAAGCCTCTAGGGAA |
| ***Peaxi162Scf03779g00019*** | GGTCTAAGCAAGAAGACCAGAA | TACCGCAACGAAGCAGACC |
| ***Peaxi162Scf00102g01226*** | GGCCTTCTTACTGAAGCTGAA | TCAGTTCTTCCTGGTAATCTTGC |
| ***Peaxi162Scf00452g00412*** | GGCCACTATTGTCAAATTGCATCA | TTAATCTCGTTATCGGTTCTTCCAG |
| ***Peaxi162Scf00016g00625*** | GCTGAACGAGTAAGAAGGGAAA | TGGAGAGACTGGACATAGTTGA |
| ***Peaxi162Scf00240g01429*** | GAGTGGCCAAGAAGCAAGTA | CCTCATCGAAATCATCATCCTCA |
| **EMSA probes** | | |
| TCH4 | GAAAATTAGGTAAAAAATTTGTG | CAACTTCAAACTTTTGCAAA |
| IRX15L | GGAGATTTAATCTCTTCACT | TGCTAGTATGTTTATAAGGAG |
| SHOU4 | CATCTAAAACTACCACATAAAT | ATTTTGCAATGAAAGTGATG |
| PILS | AATCAGTACAAGACATCAA | TTGATTTATTCCTTTGTGAG |
| IAA14 | GATGGAGATAGATAAATAACAG | GCTCATGAAGAAATAGTCTG |
| IAA16 | CTAGTGTTTGGATGAGACT | CCCACGAGAATTGAAC |
